# β-Amyloid impairs Proteasome structure and function. Proteasome activation mitigates amyloid induced toxicity and cognitive deficits

**DOI:** 10.1101/2024.10.23.619877

**Authors:** Kanisa Davidson, Mehar Bano, Danitra Parker, Karl Rodriguez, Pawel Osmulski, Maria Gaczynska, Andrew M. Pickering

## Abstract

**Background:** Impaired proteasome activity is a robust and reproducible feature in Alzheimer’s disease (AD) patients and animal models. Although amyloid beta (Aβ) may inhibit proteasomes, it is unclear if this reflects direct structural interference or indirect toxicity. The proteasome controls a myriad of critical neuronal processes with its disruption potentially driving downstream AD pathology.

**Methods:** We examined the direct impact of distinct Aβ forms on proteasome structure, dynamics, and function by evaluating enzyme activity and morphology by atomic force microscopy (AFM) and native-gel electrophoresis. To prevent Aβ-induced impairments in vitro and in AD animal models, we used allosteric proteasome activating peptidomimetics. We characterized their interactions with proteasome by AI enhanced in silico docking.

**Results:** Toxic soluble Aβ42 oligomers selectively inhibit 20S and 26S proteasome activity, while monomeric and fibrillar forms and Aβ40 show minimal inhibitory effects. Morphometric analysis of AFM images suggests Aβ42 oligomers bind laterally to the 20S proteasomes, and impair proteasome dynamics by disabling gate opening, critical for catalytic activity. Oligomer-decorated proteasomes are locked in catalytically inert cycling between closed and intermediate gate conformations. Oligomeric Aβ42 also induces disassembly of the 26S complex, increasing content of free 20S cores. Importantly, we show proteasome dysfunction induced by oligomeric Aβ42 is both preventable and reversible by employing an allosteric proteasome activator TAT-DEN, promoting gate opening and restoring enzymatic activity with the full-cycle of conformational transitions in oligomers-treated proteasomes. In cellular and animal models, restoring proteasome function via TAT-DEN mitigated Aβ-induced cell death, spatial learning/memory deficits, and survival impairments, even under conditions of pre-established Aβ deficits.

**Conclusions:** Oligomeric Aβ42 decorates the lateral sides of the 20S proteasome restricting cycling between conformations and blocking gate opening. It also reduces the pool of 26S proteasomes necessary for the ubiquitin-dependent protein degradation. We demonstrate proteasome dysfunction can be prevented and reversed by proteasome agonist that lowers 20S affinity to oligomers and promotes allosteric gate opening. We show that proteasome activation with TAT-DEN prevents cognitive deficits in AD animal models. This suggests proteasome dysfunction is a key downstream target in AD pathology and supports proteasome activation as a therapeutic strategy to restore neuronal function in AD.

## INTRODUCTION

Alzheimer’s disease (AD) is the most common cause of dementia and a leading contributor to age-related cognitive decline. Although therapies targeting β-amyloid (Aβ) have shown some efficacy, they have largely failed to halt disease progression, underscoring the need to uncover additional molecular drivers of neurodegeneration. One emerging mechanism is proteostasis failure, particularly dysfunction of the proteasome, the cell’s principal machinery for targeted protein degradation.

Reduced proteasome activity is consistently observed in AD, especially within brain regions most vulnerable to pathology such as the hippocampus and temporal cortex. In both human post-mortem tissue and AD model organisms, 20S and 26S proteasome activity is markedly diminished [1, 5–8] and correlated both with Aβ [1] and tau pathology [9]. Prior studies suggest that oligomerized or aggregated proteins, including Aβ and hyperphosphorylated tau may inhibit proteasome function [7, 8, 10], but whether this results from direct structural interference or indirect toxicity is unclear. Aβ has been proposed to act as an allosteric inhibitor of the 20S proteasome, restricting substrate access by altering gate dynamics. However, this process is under investigated [11].

The proteasome plays a central role in neuronal homeostasis, not only by degrading misfolded proteins but also by regulating plasticity-related signaling pathways. In dendritic spines, localized proteasome activity facilitates the degradation of CREB repressors and synaptic inhibitors, thereby supporting long-term potentiation, spine remodeling, and memory formation. Proteasomes are dynamically recruited to synaptic compartments via CaMKIIα and NMDA receptor signaling, linking proteolytic activity to neuronal excitation [12–20]. Disruption of these processes, whether through impaired proteasome activity or mislocalization, contributes to key phenotypes observed in AD, including dendritic spine loss, impaired synaptic signalling, and cognitive impairment [21–26].

Building on these findings, we previously reported that enhancing proteasome function can confer neuroprotection in AD models. Overexpression of the PSMB5 (β5) catalytic subunit or treatment with activators - peptidomimetics TAT1-8,9TOD and TAT1-DEN (later referred to as TAT-TOD and TAT-DEN) derived from cell-penetrating peptides (CPP) significantly increases proteasome activity and reduces Aβ-induced pathology in Drosophila, cultured neurons, and mouse models [1]. The peptidomimetics restore protein degradation capacity, attenuate synaptic dysfunction, and improve behavioral outcomes, highlighting the proteasome as a viable therapeutic target [1, 27]. Other groups have made similar findings in *C. elegans* models of AD with overexpressed β5 [28]. Comparable results have also been obtained in animal models of AD using nonspecific or indirect pharmacological interventions that included treatments with dietary supplements or marine metabolites, and elevating cAMP levels [29–31].

Despite growing recognition that proteasome dysfunction represents a key driver of AD pathogenesis, mechanism of this impairment is poorly understood. In this study, we dissect how structurally distinct forms of Aβ42 influence proteasome function, revealing oligomeric Aβ42 as a potent and conformation-selective disruptor of proteasome activity. We show that Aβ42 oligomers bind laterally to the 20S proteasome, prevent gate opening and thus disrupt the cycle of conformational transitions critical for its catalytic activity. Moreover, oligomers promote disassembly of the 26S complex, seeding proteostasis collapse. Importantly, targeted proteasome activation prevents these deleterious molecular effects of oligomeric Aβ42 and also rescues Aβ-induced toxicity in cell culture, flies, and transgenic AD mouse models. These findings provide a novel model of proteasome interactions with Aβ42 oligomers and identify proteasome activation as a viable therapeutic strategy to counteract downstream neurodegeneration.

## RESULTS

### 1. Oligomeric Aβ42 alters 20S proteasome activity

Proteasome dysfunction is a robust feature of AD, observed consistently in both post-mortem human brain tissue and transgenic animal models [1, 5–8]. This impairment has been largely attributed to direct interactions between the proteasome and pathological aggregates such as Aβ and hyperphosphorylated tau, resulting in enzymatic inhibition and a breakdown of proteostasis [7, 8, 10]. In support of this, we observed a progressive decline in proteasome chymotrypsin-like activity in hippocampal samples from AD patients that tracks with Braak staging as a marker of disease severity / progression (**Fig. 1A**).

**Fig. 1.**
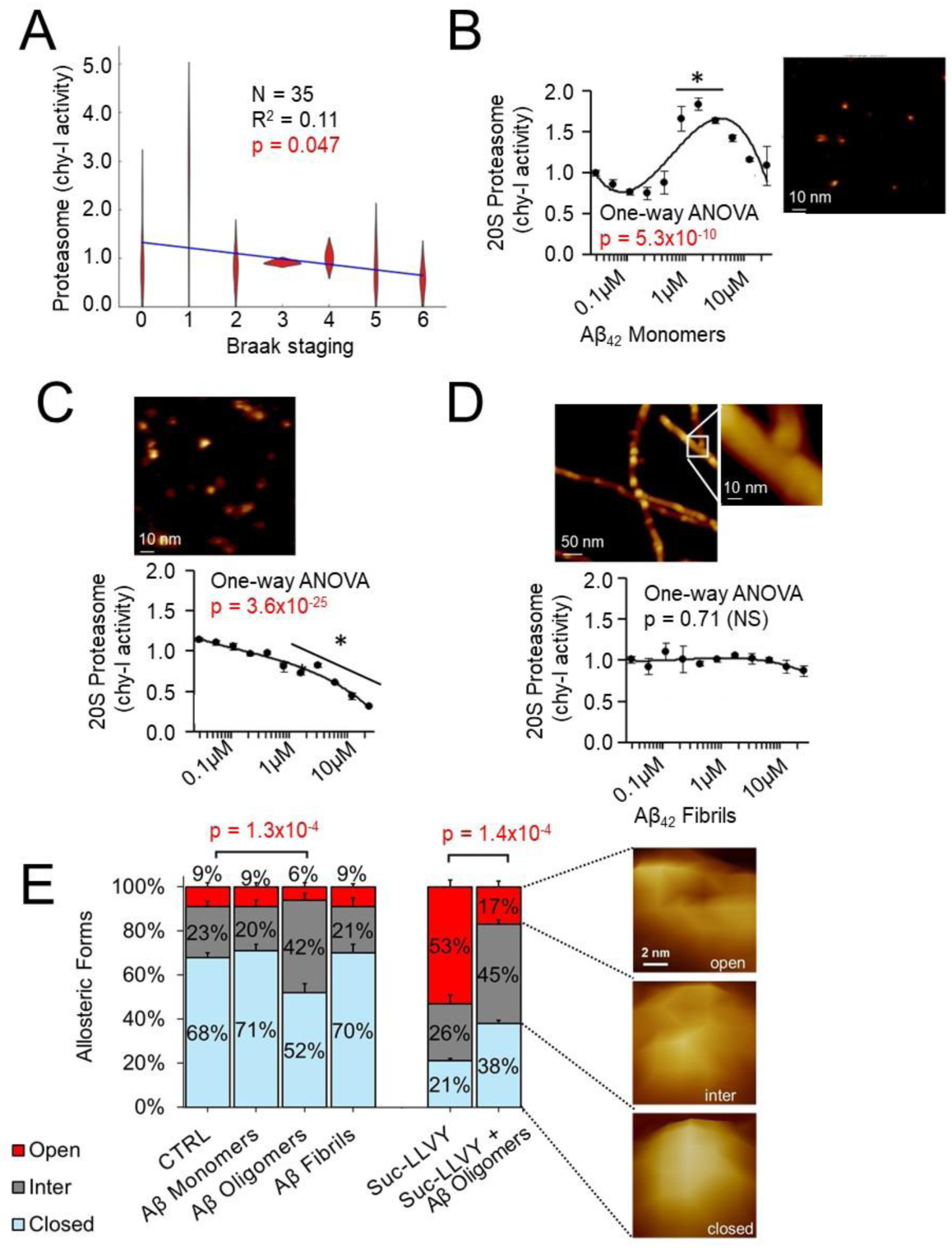
Oligomeric Aβ42 impairs 20S proteasome activity and gate opening. **(A)** Chymotrypsin-like peptidase activity (Suc-LLVY-AMC) of Alzheimer’s disease patients post-mortem human hippocampal tissue. Data represents reanalysis from our prior publication [1], stratified by Braak stage (N=35 total: Braak 0, N=7; Braak 1, N=4; Braak 2, N=7; Braak 3, N=3; Braak 4, N=2; Braak 5, N=7; Braak 6, N=5). All samples were from male patients aged 50–80. **(B–D)** Chymotrypsin-like activity of purified human 20S proteasome after incubation with **(B)** monomeric, **(C)** oligomeric, or **(D)** fibrillar Aβ42. N=4 per group. Insets show representative AFM images of Aβ particles. **(E)** Quantification of the distribution of gate conformations in the presence or absence of 2 μM oligomeric Aβ42 and 100 μM Suc-LLVY-AMC. A total of 733 particles were analyzed for vehicle-treated controls, 843 for Aβ42-treated samples, 280 for substrate-treated and 404 for Aβ42-treated samples, respectively. Statistical comparison for B-D by was performed by one-way ANOVA; asterisks indicate significance by Tukey’s post hoc test (p<0.05) with adjustment for multiple comparisons. Comparison of gate conformers’ distribution in panel E was performed using Chi-square test. Representative AFM images of zoomed-in top-view proteasome particles (pseudo-3D rendering) classified as open- intermediate- and closed-gate conformers are presented on the right.

We examined *in vitro* how different structural forms of Aβ impact proteasome function, focusing initially on the 20S proteasome, representing the catalytic core alone without UPS regulatory cap. This form degrades proteins in an ATP/Ubiquitin independent manner [32]. We started with Aβ42 peptide which is regarded as the most relevant for AD pathogenesis [33]. Following other studies, monomeric Aβ42 was prepared though sonication and centrifugation of the synthetic peptide sample to remove spontaneously formed larger forms (**Fig. 1B, inset**). Oligomers were grown via incubation of the monomers at 4°C for 24 hr under agitation, followed by centrifugation to remove larger aggregates (**Fig. 1C, inset**). Fibrils were grown through incubation of the monomers at 37°C for 24 hr (**Fig. 1D, inset**) [34]. The identity of Aβ42 forms was confirmed via Atomic Force Microscopy (AFM) imaging (**Fig. 1B-D, insets**). We assessed the influence of monomeric, oligomeric, and fibrillar Aβ42 on the proteasome function by incubation of purified human housekeeping 20S proteasomes with increasing concentrations of the Aβ42 species and testing “workhorse” chymotrypsin-like activity with a model peptide substrate (**Fig. 1B-D**). We found oligomeric Aβ to induce a potent, dose-dependent inhibition of proteasome activity (**Fig. 1C**, 1-way ANOVA p = 3.6 × 10⁻²⁵). In contrast, fibrillar Aβ42 had no measurable impact (**Fig. 1D**, 1-way ANOVA p = 0.71). Intriguingly, monomeric Aβ42 produced a modest and transient enhancement of activity at high concentrations (500 nM – 2 µM) (**Fig. 1B**, 1-way ANOVA p = 5.3 × 10^⁻10^). This apparent mild ‘activatory’ impact of high dose Aβ42 monomers is surprising and a topic for further investigation.

We extended this analysis to Aβ40, which exhibited significantly less impact on proteasome function than Aβ42. Monomeric Aβ40 impaired activity only at the highest dose tested (20 µM), while oligomeric Aβ40 produced a slight increase in proteasome activity. (1-way ANOVA p = 0.043) (**Supplemental Fig. 1**).

Taken together, these data identify oligomeric Aβ42 as the most potent and pathologically relevant species in modulating proteasome function. Its unique inhibitory profile, absent in either monomeric, fibrillar or Aβ40 forms, supports a model in which soluble Aβ42 oligomers engage with the 20S proteasome through distinct, conformation-sensitive mechanisms. Accordingly, all subsequent experiments in this study focus specifically on oligomeric Aβ42.

### 2. Oligomeric Aβ42 affects partition of 20S proteasome gate conformers compromising the ability to accept substrates

To dissect how oligomeric Aβ42 modulates 20S proteasome conformation, we employed high-resolution AFM imaging to visualize single proteasome particles in a top-view (standing) orientation, exposing their α-rings for structural assessment, as in our previous works [27]. Particles were classified into three discrete gate conformations: closed, intermediate, and open (**Fig. 1E** **inset**). Incubation with oligomeric Aβ42 significantly shifted this distribution reducing the contribution of closed-gate and open-gate conformers, while increasing the contribution of intermediate state (**Fig. 1E**, Chi-square p = 1.39×10^-4^). Addition of a model peptide substrate for proteasomal chymotrypsin-like peptidase (Suc-LLVY-AMC) to the control proteasomes led to a massive increase in the content of open-gate conformers, at the expense of closed-gate forms (**Fig. 1E**), consistent with the model of allosteric gate opening of the 20S proteasome by a substrate [35]. The increase in content of the open form after the addition of the substrate was detectable in the presence of Aβ42 oligomers, however it was much less pronounced. Control proteasomes displayed a shift from 9% open to 53% open with addition of substrate. In the presence of Aβ oligomers this was reduced to a shift of 6% open to 17% open. These findings suggest compromised substrate entry in the presence of Aβ42 oligomers (**Fig. 1E**). Interestingly, the proportion of intermediate forms remained nearly unchanged in the presence of oligomers, whether or not the substrate was added (42% vs 45%, respectively; **Fig. 1E**). In contrast, we did not observe robust changes in gate dynamics in the presence of Aβ42 monomers or fibrils (**Fig. 1E**).

These findings indicate that the Aβ42 oligomers productively interact with the 20S core proteasome. Therefore, we next setup to explore potential binding areas for the inhibitory ligands.

### 3. Oligomeric Aβ42 preferentially binds to the lateral surface of closed-gate and intermediate-gate conformers

Morphometric analysis of AFM images of single 20S proteasome particles allowed us to characterize the specific shape and dimensions of hundreds of top-view molecules. Under applied conditions the surface of proteasome α ring (“α face”) to the practical depth of less than 3 nm is accessible for imaging. The complete image of the top (**Fig. 1E** **inset**) is a composite of up to 6 scanning lines, each line with up to 6 pixels covering the α face. It takes up to 2 seconds to scan the full α face. Scanning the center horizontal line used to distinguish the gate conformers (see Materials and Methods) takes about 2 milliseconds.

Based on statistically significant differences between control and Aβ42 treated proteasomes we choose for analysis the following morphometric parameters of a single particle: Aspect – the ratio between maximal and minimal diameters measured from the particle’s center of gravity (***Fig. 2*** ***Left***), X-extend - the length of center line which serves as the approximation of a real-time diameter of the α face with an assigned gate conformation (***Fig. 2*** ***Middle***), Z Max - the maximal height of the particle (***Fig. 2. Right***), Perimeter – the total length of the projected contour (***Supplemental Fig. 2 Left***), and Area – the projected footprint (***Supplemental Fig. 2. Right***). X-extend (diameter, ***Fig. 2. Left***) is the sole parameter corresponding precisely to the assigned conformation, while other parameters combine the features of the X-extend assigned conformer and the ensemble. Since the conformers are morphologically distinct, we pairwise compared selected parameters of control with Aβ oligomer-treated proteasomes. It is worth noting that the morphometric differences between allosteric forms in control particles, evident in ***Supplemental Table 1***, match those detected by cryoEM imaging. For example, the area increases from closed through intermediate to open forms (***Supplemental Table 1***). Accordingly, the solvent-accessible surface of α rings in cryoEM-identified forms increases by about 4% from closed to intermediate, and by 5% from closed to open [3], see Materials & Methods for details).

**Fig. 2.**
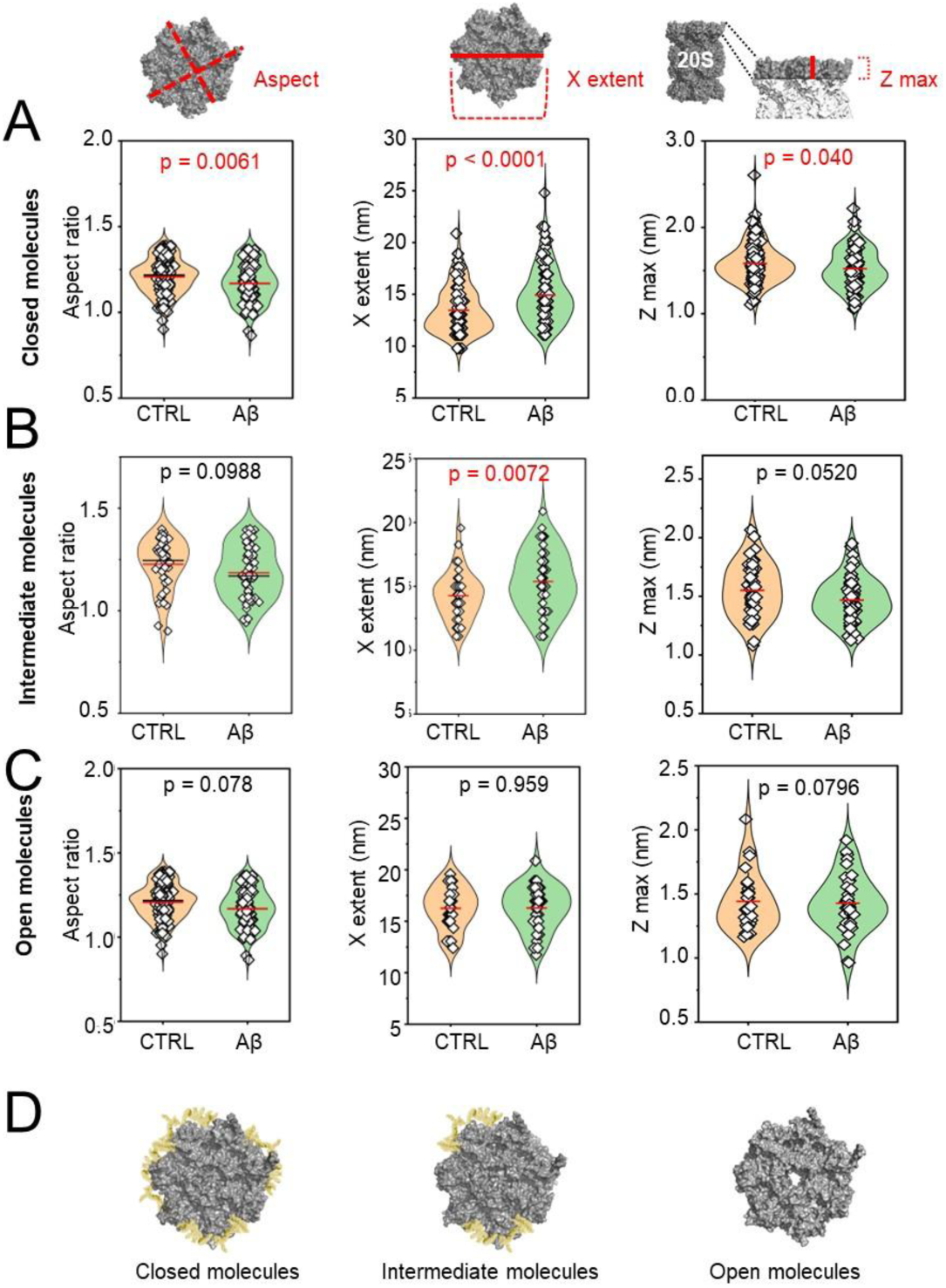
Treatment with oligomeric Aβ42 induces changes in shape of closed- and intermediate- but not open-gate 20S proteasomes. Morphometric analysis of AFM-imaged top-view 20S proteasomes classified into three conformational states: closed, intermediate, and open-gate, was performed for proteasomes pre-incubated with the vehicle (control) or 2 μM oligomeric Aβ42. The following morphometric parameters were assessed: Aspect (***Left***) – the ratio between maximal and minimal diameters measured from the particle’s center of gravity, X-extend (***Middle***) - the length of center line which serves as the approximation of a real-time diameter of the α face with an assigned gate conformation; the features of center line are actually used to determine the gate status (see Materials and Methods), Z max (***Right***) - the maximal height of the particle. The rows represent conformers identified as: **(A)** closed-gate, **(B)** intermediate, **(C)** open-gate. **(D)** Proposed model with closed conformers showing the lateral coating with Aβ42 oligomers, intermediate conformers showing evidence of Aβ42 coating but with less pronounced effects and a higher morphometric variance. Open conformers appear resistant to coating by Aβ42 oligomers. Each point represents a single proteasome particle. Control group included 134 closed, 45 intermediate, and 30 open gate particles; Aβ42-treated group included 110 closed, 60 intermediate, and 38 open gate particles. Red horizontal lines represent averages. Statistical analysis was conducted via pairwise comparison; p < 0.05 was considered significant (colored red). Molecular models of human 20S proteasomes were rendered from pdb 6MSB (closed), 6MSH (inter) and 6MSK (open) [3].

As evident in **Fig. 2A** and ***Supplemental Fig. 2A***, closed-gate particles treated with Aβ42 oligomers appeared larger than control, with significantly larger X-extend (real-time diameter), area and perimeter. However, two parameters defied the trend, with oligomers treatment resulting in significantly lower Aspect ratio and lower Z (Height) max (**Fig. 2A** and ***Supplemental Fig. 2A***). Lower Aspect indicates a more rounded, circle-like shape. The larger X-extend, Area, and Perimeter suggest presence of an additional mass, such as oligomers, to the lateral surface. In turn, a lower Z max (maximal height) in the treated proteasomes does not support a notion of Aβ oligomers binding at the top of α face, when a higher rather than lower Z max would be expected (**Fig. 2A, *Supplemental Table 1***). Consistently, top binding is not supported by “real time” data collected along the 10 nm of center scan line, where both average real time Z (height 1.32 nm vs 1.31 nm; control vs oligo-treated, p=0.53) and real time maximal Z (height 1.60 nm vs 1.54 nm; p=0.2) did not differ significantly between control and oligo-treated particles, instead showing additional mass at the rim of α face (***Supplemental Fig. 3***). Overall morphometric analysis of closed conformers suggests lateral not top binding of Aβ42 oligomers.

Similar to the closed conformers, morphometric analysis of intermediate forms suggests a possibility of lateral binding of oligomers as well, with a significant enlargement of the real time diameter (X-extend). Other parameters of the intermediate particles showed trends similar to the closed forms, however without reaching statistical significance (**Fig. 2B**: lower Aspect ratio and Z max, ***Supplemental Fig. 2B***: higher Perimeter and Area, see also ***Supplemental Table 1***). This suggests intermediate conformers to be less susceptible to coating than fully closed conformers, but may also indicate that morphological features of intermediate particles are less permissive for detecting the binding.

Contrary to the closed conformers, none of the morphometric parameters of the particles classified as open differed significantly between control and Aβ42 oligomers – treated particles (**Fig. 2C** **and *Supplemental Fig. 2C***). Accordingly, we hypothesize that open conformers, unlike closed and intermediate, are much less receptive to coating with oligomers (**Fig. 2D**).

Collectively, these findings demonstrate that interactions of the 20S proteasome particles with Aβ42 oligomers are spatially restricted to the *lateral surface* of closed- and intermediate-gate proteasome conformers (**Fig. 2D**). The top of the α-face is left unobstructed pointing at the allosteric restrictions rather than direct gate blocking as the culprit behind proteasome inhibition by oligomers.

### 4. Oligomeric Aβ42 drives 26S proteasome disassembly

We next analyzed the impact of oligomeric Aβ42 on the 26S proteasome, a multi-subunit complex composed of the 20S catalytic core and one or two 19S regulatory caps, which enables the degradation of polyubiquitinated proteins [36]. Similar to our studies shown in **Fig. 1B-D** with 20S proteasome, we assessed chymotrypsin-like peptidase activity of purified human 26S proteasome treated with increasing concentrations of monomeric, oligomeric or fibrillar Aβ42 in the presence of ‘MAD’ (MgCl_2_, ATP, and DTT) for 26S stabilization. We found both monomeric and oligomeric Aβ42 impaired the 26S proteasome activity in a dose dependent fashion (p=0.007 and p = 3.6×10^-16^, respectively, **Fig. 3A-C**) with oligomeric Aβ42 displaying higher efficacy. In contrast, we saw no inhibition of 26S by fibrillar Aβ42 (**Fig. 3C**). Because the α-ring is occluded by the 19S regulatory cap in 26S proteasome, it was not possible to directly assess 26S proteasome gate dynamics with AFM.

**Fig. 3.**
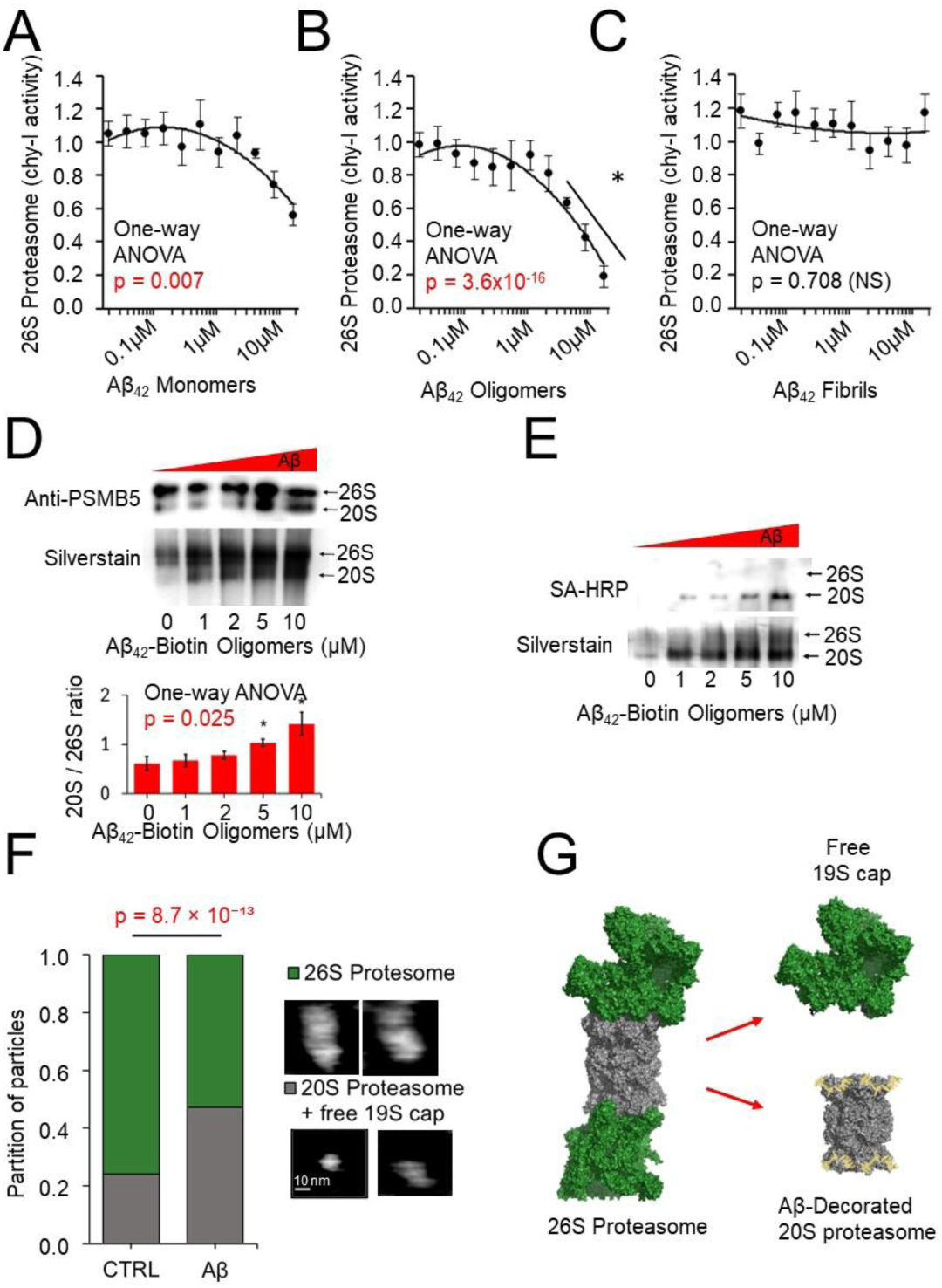
Oligomeric Aβ42 impairs 26S proteasome activity and assembly. (A–C) Chymotrypsin-like peptidase activity of purified human 26S proteasomes following incubation with **(A)** monomeric Aβ42, **(B)** oligomeric Aβ42, and **(C)** fibrillar Aβ42. N = 4 per group. **(D,E)** Native PAGE immunoblot of 26S proteasomes incubated with increasing concentrations of (**D**) oligomeric Aβ42 and **(E)** biotinylated- oligomeric Aβ42. In D. Blots were probed for the β5 subunit, with corresponding silver stain shown. Arrows indicate positions of intact 26S and free 20S proteasomes. In E. Biotin-strepdavadin conjugation was performed with streptavidin-HRP followed by detection. Equivalent amounts of 26S proteasome were loaded per lane; variations in silver staining are attributed to amyloid decoration rather than differences in protein loading. Quantification shows 20S-to-26S proteasome ratios from native gels N = 4 per condition. **(F)** Morphometric analysis of AFM images of purified human 26S proteasome imaged in the absence and presence of 2μM Aβ42 oligomers. Distinct assemblies of the proteasome were classified based on their Length, as demonstrated in **Supplemental Fig. 4**. Results show the partition of 19S-decorated core proteasomes (single or double capped) to free 20S cores and 19S regulatory particles. Participation of the latter significantly increased in the oligomers-treated samples, suggesting disassembly of the 26S complexes. Raw images of representative particles are presented on the right. Analysis included 536 control particles and 382 Aβ42 oligomer treated particles. Significance of partition difference was assessed with the Chi-square test. See **Supplemental Fig. 4**. for the length distribution and morphometric properties of particles. **(G)** Model of oligomers-supported loss of integrity of the 26S proteasome (molecular models rendered from pdb 5t0c; [4]. Statistical significance In A-D determined by one-way ANOVA; and Chi-square in E. asterisks indicate p < 0.05 by Tukey’s post hoc test adjusted for multiple comparisons.

The 20S and 26S proteasome exist in a dynamic state with the 19S cap capable of detaching and reattaching [37]. We assessed the relative composition of 20S and 26S proteasomes in the presence of oligomeric Aβ42 via Native PAGE followed by immunoblotting against a core 20S proteasome subunit (β5; PSMB5). In this approach we can visualize distinct bands for the 20S and 26S proteasomes (**Fig. 3D, arrows).** Silver staining additionally discerns separate bands for double-capped and single-capped 26S proteasomes, merged in lower-resolution immunostaining, likely due protein diffusion during transfer to a PVDF membrane. We found that incubation with oligomeric Aβ42 led to a dose-dependent reduction in intact 26S complexes and a concomitant increase in free 20S cores (**Fig. 3D**). These findings suggest that Aβ42 oligomers promote disassembly of the 26S proteasome. We ran a parallel experiment using biotinylated-Aβ42 oligomers, where we observed a distinct band of biotinylated-Aβ42 aligning with 20S proteasome; interestingly we did not observe a band corresponding to the 26S proteasome assembly. This suggests that oligomeric Aβ42 can interact only with the 20S proteasome, but when bound to 20S it prevents re-formation of 26S proteasome assembly (**Fig. 3E**). We hypothesize that oligomeric Aβ42 prevents interactions between the 20S core and 19S regulatory particles. These actions result in the inhibition of the 26S activity both by shifting the equilibrium toward free 20S core and 19S caps, and by direct inhibition of the core.

As an independent verification of our findings, we employed AFM imaging of the 26S proteasomes followed by particle analysis. Particles were grouped by size (Length; maximal length of a particle) in [1] 20S proteasomes and free 19S regulatory particles, [2] single of double caped 26S proteasomes. In the absence of Aβ we found 75.8% of our sample to consist of 26S proteasomes, this was reduced to 52.9% in the presence of Aβ oligomers (**Fig. 3F**, ***Supplemental Fig. 4*,** p = 8.7 × 10⁻¹³). Based on these findings we propose that while Aβ42 oligomers have a low binding affinity to the capped 20S core, once bound they prevent the core-19S interactions and thus shift the equilibrium toward more abundant free 20S particles (**Model** **Fig. 3G**).

### 5. Proteasome allosteric activation mitigates Aβ42 induced proteasome inhibition by shifting conformational equilibrium toward oligomers-refractory open-gate forms

We sought to test if pharmacological manipulation to alter proteasome conformational dynamics could reduce Aβ inflicted proteasome dysfunction. To meet this goal we have developed proteasome peptidomimetic activators TAT1-8,9TOD (TAT-TOD) and TAT1-Dendrite (TAT-DEN). The compounds activate the 20S and 26S proteasome *in vitro*, *in cellulo* and *in vivo,* pass the BBB and show no toxicity in animal models, with a detailed characterization provided in our prior publications [1, 27]. Because of its more favourable drug-like properties (*in vitro* kinetics, stability, detectability in mouse brain; [1]), we focused on TAT-DEN (**Fig. 4A**).

**Fig. 4.**
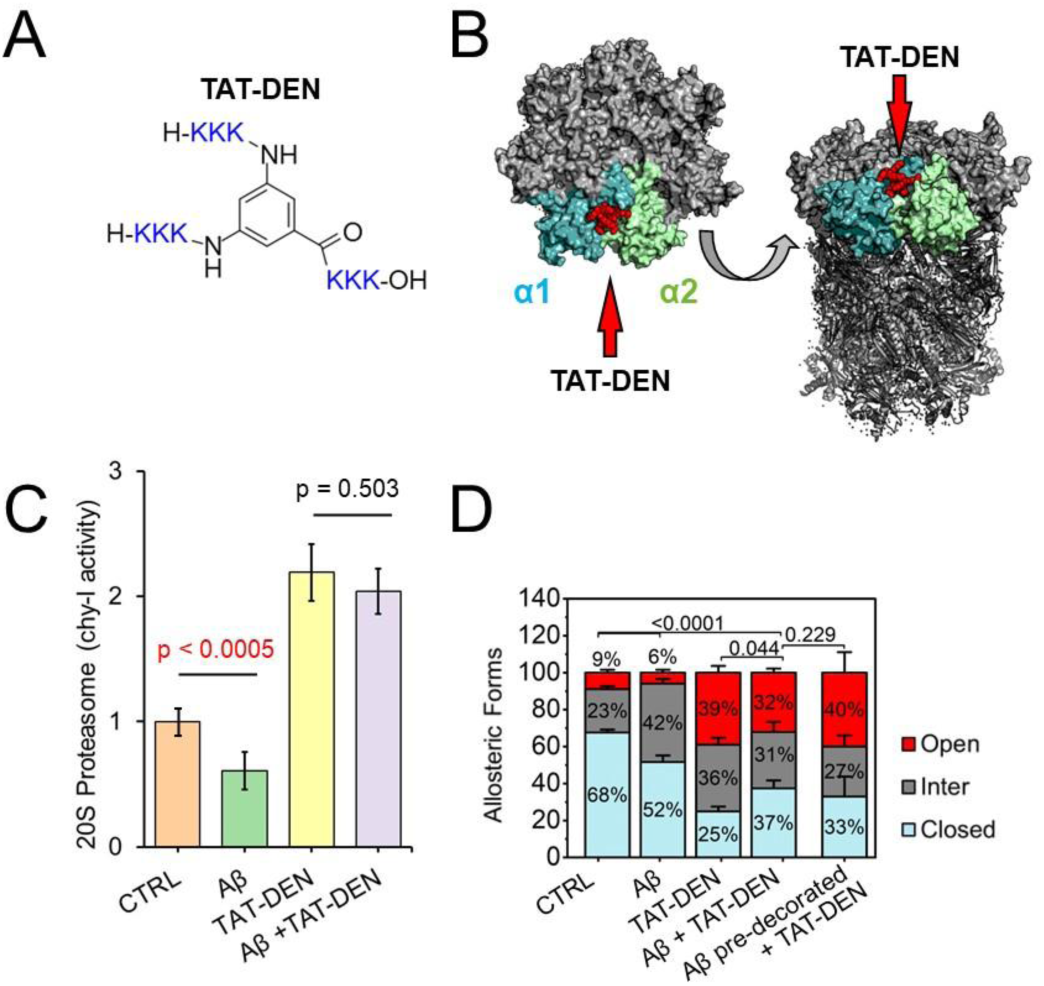
Proteasome agonist TAT-DEN prevents Aβ-induced proteasome impairment. (A) Structure of proteasome agonist TAT-DEN. **(B)** Molecular docking simulation of TAT-DEN poses. Docking of TAT-DEN: Surface rendering of top and side views of the 20S, α1/α2 subunits colored. Human housekeeping 20S structure model used for docking: pdb 5LE5, [2]. **(C)** Chymotrypsin-like activity of purified human 20S proteasome ± 2μM Aβ42 oligomers ± 1μM TAT-DEN. N=3. **(D)** AFM analysis of purified human 20S proteasome ± 2μM Aβ42 oligomers ± 1μM TAT-DEN. Proteasomes were premixed and preincubated with ligands (or vehicles) for 1 hr RT before AFM imaging (CTRL, Aβ, TAT-DEN, Aβ + TAT-DEN) or TAT-DEN was added to the proteasomes already immobilized on mica and pre-treated for 30 min with the Aβ oligomers. Particles were classified into closed, intermediate, and open gate conformers. The number of particles analyzed: 733 (vehicle control), 843 (with oligo Aβ42), 270 (with 1 μM TAT1-DEN), 171 (Aβ42 + TAT1-DEN; “mix”), and 148 (Aβ pre-decorated + TAT-DEN). P-values based on Student’s t-test except D which employs FDR corrected Chi-square.

Using AI-enhanced molecular dynamics simulations and *in silico* docking (see Materials & Methods) we identified an allosteric proteasome binding pocket for TAT-DEN between α1 and α2 core subunits (**Fig. 4B**). Recapitulating our prior work, we show TAT-DEN to enhance activity of purified 20S as well as 26S proteasome *in vitro (****Supplemental Fig. 5***) [1, 27]. Importantly, we show that TAT-DEN is still able to enhance proteasome function in the presence of oligomeric Aβ42, reversing the proteasome inhibition by Aβ (**Fig. 4C**). Utilizing AFM imaging we found that treatment with TAT-DEN shifts the equilibrium of conformers toward the majority of open-gate (39%) and intermediate (36%) forms, leaving the closed forms in a minority (25%; **Fig. 4D**). Notably, proteasomes treated with oligomeric Aβ42 and TAT-DEN showed a markedly similar distribution of conformers to those treated with TAT-DEN alone. This observation was valid regardless of the order of addition of oligomers and TAT-DEN, with no significant differences in particles partition (**Fig. 4D**). We found, treatment with TAT-DEN prevents Aβ42 induced shift toward the excess of intermediate conformers and supports formation of open gate proteasomes essential for accepting substrates (**Fig. 4D**).

### 6. Treatment with TAT-DEN prevents binding of oligomeric Aβ42 to intermediate-gate proteasome conformers

With the prominent rescue effects of TAT-DEN on both the catalytic activity and partition of allosteric forms of Aβ42 oligomers - treated 20S proteasomes, we examined the putative status of Aβ oligomers binding to the 20S particles in the presence of TAT-DEN. Similarly to experiments in ***Fig. 2***, control and oligomers-treated particles, we compared morphometric parameters of proteasomes treated with TAT-DEN alone and in combination with Aβ oligomers. We noticed that the particles identified as closed presented significantly lower Aspect and Z max, but higher X-extend, Perimeter, and Area (**Fig. 5A, *Supplemental Fig. 6B and Supplemental Table 2***). These features were recognized as Aβ oligomer-coating similar to proteasomes treated with Aβ oligomers alone (**Fig. 2A**). However, when both the ligands were present, intermediate conformers did not display the increased X-extend expected when proteasomes are coated with oligomers (**Figs. 2B**, vs **5B**). Likewise other morphological parameters (aspect, Z max, perimeter, area) were not altered between TAT-DEN and TAT-DEN-Aβ-oligomers – treated intermediate proteasomes (**Fig. 5B, *Supplemental Fig. 6B and Supplemental Table 2***). Likewise, TAT-DEN and TAT-DEN-Aβ-oligomers – treated open proteasomes did not show a significant enlargement of X-extend, perimeter or area, or lowering of Z max and aspect, that would suggest coating with oligomers (**Fig. 5C, Supplemental Fig. 6C and Supplemental Table 2**). This is similar to our findings of no putative coating in Aβ-oligomers – treated open proteasomes shown earlier (**Fig. 2C** **and Supplemental Fig. 2C**). We hypothesize that TAT-DEN prevents binding of oligomers to the intermediate forms and even induces a release of (shakes-off) the already bound oligomers thus leaving the intermediate forms free to open the gate, as was shown in **Fig. 4D**.

**Fig. 1.**
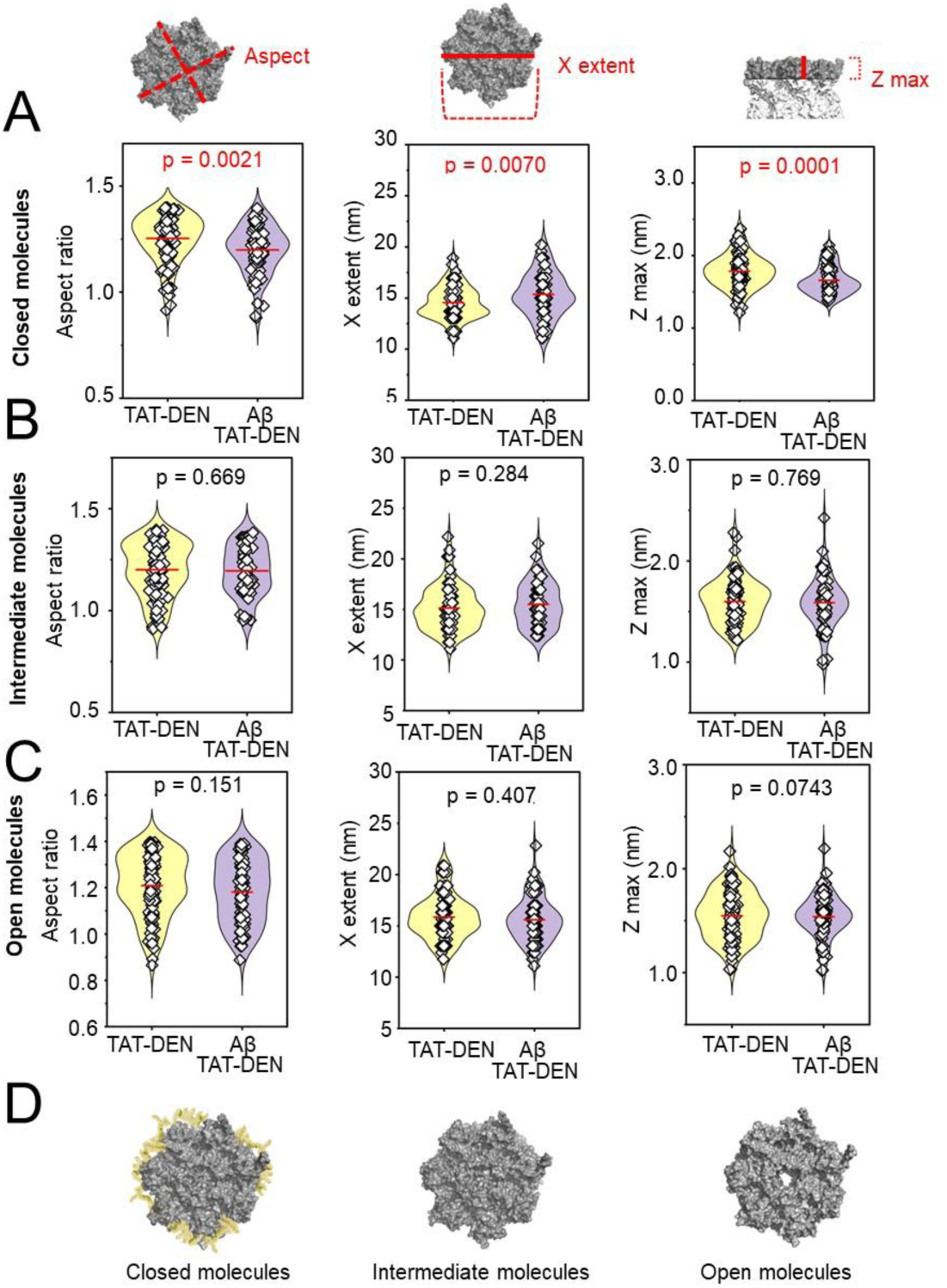
Morphometric shifts induced by TAT-DEN in 20S proteasome treated with oligomeric Aβ42 are limited to closed-gate conformers. Morphometric analysis of AFM-imaged top-view 20S proteasome particles was performed as in Fig. 2, but in the presence of 1μM TAT-DEN. Three morphometric parameters: Aspect (***Left***), X-extend (real time diameter; ***Middle***) and Z max (***Right***) were analyzed for particles identified as closed- **(A)**, intermediate- **(B)** and open-gate conformers **(C)**, as before. **(D)** In the proposed model closed conformers are laterally coated with Aβ oligomers even in the presence of TAT-DEN. However, open and intermediate conformers appear resistant to the coating by Aβ oligomers. Each point represents a single proteasome particle. TAT-DEN group included 80 closed, 62 intermediate, and 106 open gate particles; TAT-DEN Aβ42-treated group included 91 closed, 92 intermediate, and 82 open gate particles. Statistical analysis was conducted via pairwise comparison; p < 0.05 was considered significant (colored red).

With the intermediate forms apparently liberated from oligomers by TAT-DEN we asked how closed conformers are impacted by the activator ligand. For that we analysed the Aspect values (a measure of the shape “roundness”) of free and ligand bound closed particles. In bulk, the Aβ oligomer-treated closed particles were significantly more round than controls, showing a decreased Aspect (**Fig. 2A** **& 5A**). Importantly, as presented in ***Supplemental Fig. 7*,** the frequency plot of aspect values for oligomers-treated particles revealed a prominent population of low-Aspect particles (peak I) not detected in the control proteasomes and likely representing the well-coated particles. Peak II (higher Aspect, less rounded particles), detected as the major population in controls, was present but much less prominent in the Aβ oligomer treated cohort. Peak III (highest Aspect, elongated particles) was well-discerned in all cases but show a minor contribution in controls and much more prominent, shifted to lower values in the oligo-treated particles. We propose that Peak III represents the dynamic particles that are in transition to intermediate conformers both in the control and treated populations, and that those particles are still oligo-coated in the latter. When a similar analysis was performed for the proteasomes treated by combination of TAT-DEN and Aβ42 oligomers, the coated-particles peak I was well discerned, however the “control” (free from Aβ42 oligos -closed) peak II was much more prevalent than in the absence of TAT-DEN. Peak III was more similar to the peak in control particles than of coated particles. Additional peaks present in the TAT-DEN – treated proteasomes likely reflected the much more robust dynamics of activated proteasomes (***Supplemental Fig. 7***). Taken together, the data suggests that closed particles have a lower affinity to the Aβ oligomer in the presence of TAT-DEN, paving the way to shaking-off the oligomers during the switch to the intermediate form.

### 7. Treatment with TAT-DEN restores the cycle of conformational transitions of the gate restricted by oligomeric Aβ42

Both Aβ oligomer-treated and TAT-DEN-treated proteasomes display a high content of intermediate-gate conformers, however they differ dramatically in their catalytic proficiencies and in partition of other gate forms (**Fig. 4** **E, F**). To further explore the conformational dynamics of proteasome particles, we attempted to temporally trace the gate transitions using AFM imaging. As described above, the classification of conformers was based on the shape of center line scanned horizontally across the top-view 20S core particles. The AFM probe scans a field of particles from left to right, then quickly returns in the same line right to left (“trace” and “retrace” images, respectively), before jumping vertically to the next line. Thus, particles that are positioned close to the right border of the field are scanned twice within a short time span, allowing for an assessment of the gate dynamics within the time-frame suitable for allosteric transitions. We approximated the time between trace and retrace scans of the same particles positioned within 0 to 130 nm from the right border as 10 to 50 milliseconds. We found that within that time frame 17% of the control particles are either open in both trace and retrace (“stable open”) or switch between intermediate and open (either direction; **Fig. 6A**). Such particles can be considered capable of accepting substrates (substrate-competent). Stable closed forms represent 58% of the total ensemble, whereas stable or changing to closed intermediates account for 26% (**Fig. 6A**). The proportion of substrate-competent particles in Aβ oligomers-treated proteasomes was lowered to less than 8%, with a striking increase in content of stable intermediate forms to 32% (**Fig. 6B**). Interestingly, the total participation of changing forms (inter to open or inter to closed) was lowered from 30% in control to less than 16% in oligomers-treated cohort (**Fig. 6B**), strongly suggesting impairment of conformational transitions by Aβ oligomers. Aβ oligomers-coated particles appear inefficient in switching toward the open conformer, leaving the majority of the proteasome molecules circling idly between closed and intermediate forms. This block was relieved by the treatment with TAT-DEN. Not surprisingly, the activity-enhancing TAT-DEN treatment of control 20S proteasomes elevated the content of substrate-competent forms to 51%, while reducing the content of closed and stable intermediate forms (**Fig. 6C**). The positive effect was retained in the presence of oligomers: substrate–competent forms accounted for 36% of all the particles. TAT-DEN also appeared to relieve the Aβ oligomers-promoted block on switching toward an open gate conformation (**Fig. 6D**). The findings are summarized in the model in **Fig. 6E**. We propose that binding TAT-DEN to the proteasome α face lowers affinity of Aβ oligomers to the closed and especially to the intermediate-gate conformers, restoring their ability to allosterically open the gate and accept substrates.

**Fig. 6.**
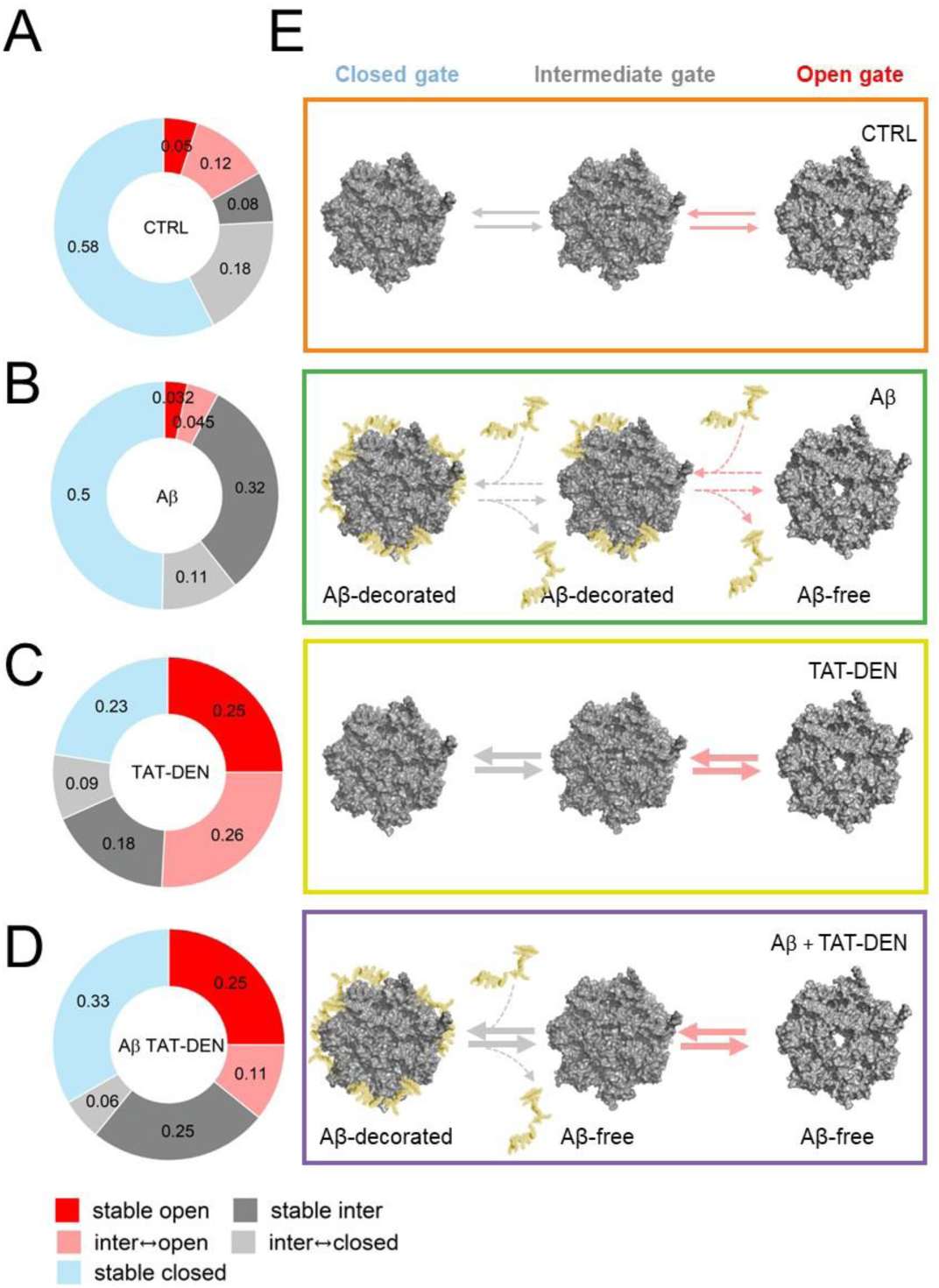
TAT-DEN restores dynamic conformational transitions of 20S proteasomes impaired by Aβ42 oligomers. (A-D) Doughnut charts show the distribution of 20S proteasome gate conformers identified by time-resolved AFM trace–retrace imaging (10–50 ms interval) across four conditions, from top to bottom: **A.** control, **B.** treated with Aβ42 oligomers, **C.** with TAT-DEN or **D.** Aβ42 and TAT-DEN. Closed (blue), intermediate (grey), open (red), and switching (light red or light grey) conformers were quantified. Treatment with Aβ42 oligomers increased the contribution of stable intermediate and closed forms while reducing substrate-competent (open and switching to/from open) particles. Treatment with TAT-DEN enhanced the proportion of open and switching forms almost restoring the gate mobility in Aβ-treated samples. **(E)** A model proposing that the Aβ42 oligomers induced “locking” the proteasomes in idle cycling between closed and intermediate forms, and TAT-DEN stimulated a shift toward substrate-competent forms, even in the presence of oligomeric Aβ42. Arrow thickness represents state transition

### 8. Proteasome inhibition increases in cellulo Aβ toxicity and proteasome activators protected against in Aβ-induced cell death

Using neuroblastoma SK-N-SH cells we demonstrate that proteasome inhibition increases susceptibility of cells to Aβ toxicity. Treatment with the proteasome selective inhibitor Mg132 does not induce cell death in itself at our experimental dose but co-treatment with oligomeric Aβ42 and Mg132 causes more pronounced cell death than treatment with oligomeric Aβ42 alone **(Fig. 7A**). We found that augmentation of proteasome function through treatment with our agonists protects against Aβ42 toxicity. Treatment of cells with our proteasome agonists TAT-DEN of TAT-TOD both reduces cell death as a consequence of Aβ42 (**Fig. 7B, Supplemental Fig. 8A**). Furthermore, we show the impacts of proteasome activators on Aβ toxicity are driven directly as a consequence of proteasome activation. Using TAT1 4,5/8,9TOD, a structurally similar peptidomimetic but with a disrupted pharmacophore and poor function as a proteasome activator [24], we show no significant protection against Aβ42 toxicity (**Supplemental Fig. 8B**). These results suggest the impacts on Aβ toxicity to stem directly from the role of our compounds as proteasome activators.

**Fig. 7.**
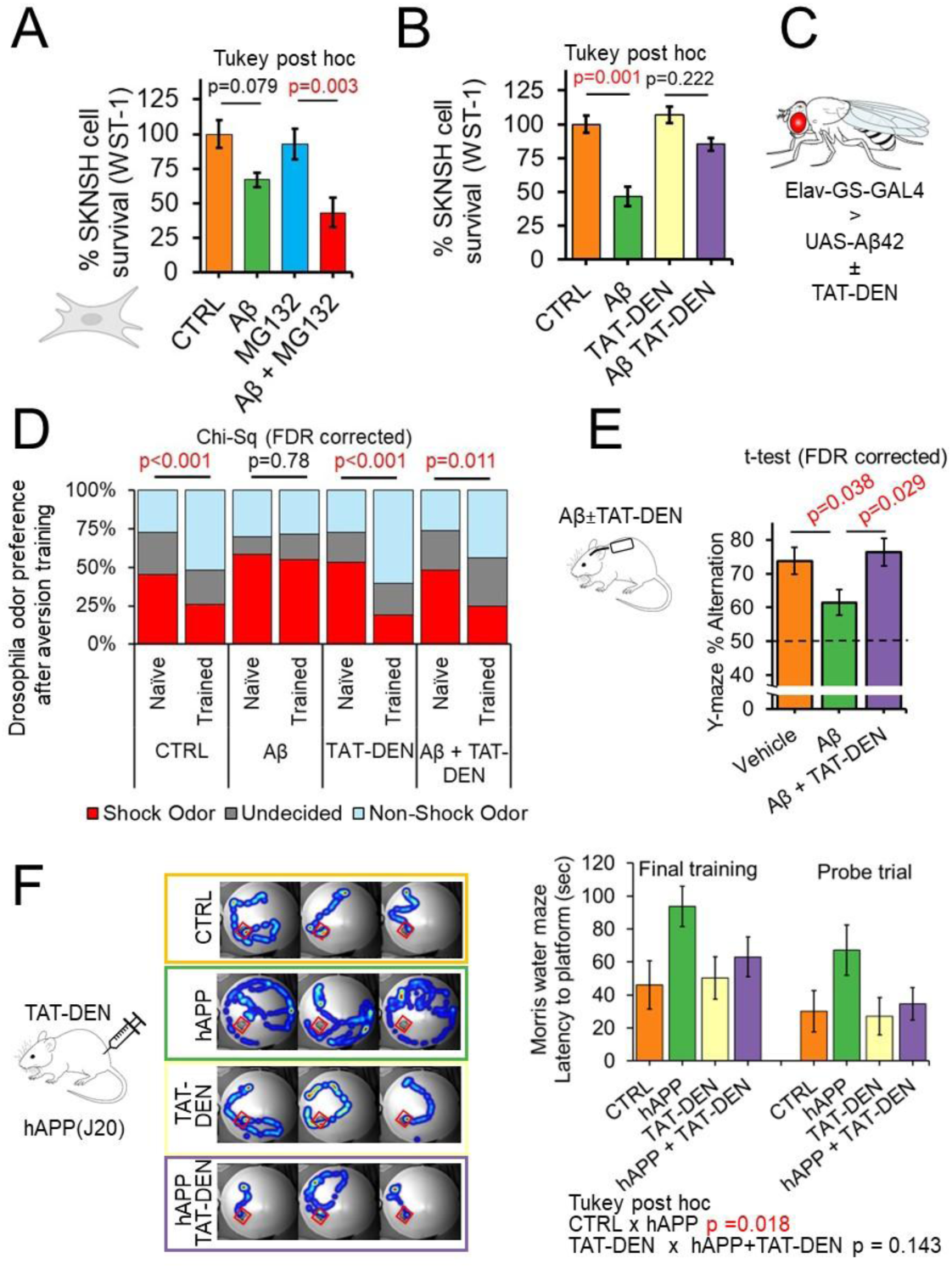
Aβ42 produces deficits in working memory, memory formation and acquisition which may be rescued by either chronic or acute treatment with proteasome agonists. (A-B) WST1, cell viability assays in SK-N-SH cells seeded at 100,000 cells per ml after 4 days treatment with 5 μM oligomeric Aβ42. Cell death was modulated under co-treatment with 0.1μM MG132 (**A**) or 1μM TAT-DEN (**B**). N = 15 wells/condition. **(C-D)** Olfaction aversion training assay in day 30 old flies comparing Elav-GS-GAL4>UAS-Aβ42 ± 200 RU486 (CNT/Aβ42 O.E.) in the absence and presence of 1μM TAT1-DEN or vehicle mixed directly into food, N=60-167/group. **(E)** Osmotic pumps surgically implanted in 3 months old C57BL6J mice to deliver to the hippocampus +/- 1.25μmol of Aβ42 +/-100pmol of TAT-DEN per hr for 21 days. Spontaneous alternation in Y-Maze assessed. N=11 animals/group. **(F)** Morris water maze in 18-month-old hAPP(J20) and littermate controls. Animals received 3, 1 minute training sessions from different locations on day 1 with a raised platform. Animals were IP injected on Day 2 with 400nmol/kg of TAT-DEN or

### 9. Proteasome activators protect against Aβ related cognitive and survival deficits in Drosophila

We next sought to determine if proteasome augmentation would protect against toxicity effects of Aβ in an animal model. We have previously reported that augmentation of proteasome function modulates the formation of Aβ through impacts on amyloid precursor machinery [1]. To determine if there are downstream protective impacts independent of Aβ synthesis we utilized a Drosophila model of Alzheimer’s disease which overexpresses pre-formed Aβ42 under control of a pan-neuronal Elav driver (Elav-GS-GAL4>UAS-Aβ42) (**Fig. 7C**). This model experiences survival and cognitive deficits. We first demonstrated treatment with our proteasome agonists TAT-DEN and TAT-TOD to have no significant impact on lifespan in wild-type (Orgeon-R) flies (**Supplemental Fig. 9A**). We found oral administration of either TAT-DEN or TAT-TOD to delay mortality as a consequence of Aβ42 overexpression (**Supplemental Fig. 9B**). We found TAT-DEN to delay only early life toxicity effects from Aβ42 while TAT1-8,9TOD to delay both early and late life toxicity effects (**Supplemental Fig. 9B**). To investigate the impacts on cognition, a key aspect of AD pathology, we employed an olfaction aversion training assay. Flies are exposed to two distinct odors while given an electric shock under exposure to one. The flies are then placed in a 2-sided chamber with the two odors released from either side. Aversion to the odor associated with an electric shock was quantified as a measure of learned response. We found control flies to show a training induced aversion to the odor associated with an electric shock. Treatment of control flies with proteasome agonists produced no effect on training induced odor avoidance (**Fig. 7D**). We show training induced avoidance of the shock-linked odor is lost in flies which overexpress Aβ42, however, treatment of flies which overexpress Aβ42 with either TAT-DEN or TAT-TOD restored training induced odor avoidance (**Fig. 7D, Supplemental Fig 9C**).

### 10. Acute and chronic treatment with proteasome activators protects against Aβ related cognitive and deficits in mice

We next investigated mammalian translational relevance, whether proteasome augmentation can protect against Aβ induced cognitive deficits in mice. To do this we employed mice fitted with an osmotic pump to deliver either Aβ42 and vehicle or Aβ42 in conjunction with TAT-DEN or TAT-TOD directly to the brain for 21 days. Treatment with Aβ42 produced deficits in working memory in mice as assessed by arm alternation in a Y-maze assay (**Fig. 7E**). We showed co-treatment with TAT-DEN to significantly improve alternation back to wild type levels. Treatment with TAT-TOD also produced improvements but unlike TAT-TOD these did not reach statistical significance (**Supplemental Fig. 10**). For this reason, we focused our subsequent investigations on TAT-DEN.

Previous studies have shown acute treatment with proteasome inhibitors to produce deficits in long term memory formation and retrieval [38–40]. Our running hypothesis is that Aβ induced inhibition of proteasome function may contribute to deficits in memory formation. We thus hypothesized that acute treatment with proteasome activators might rescue AD-like deficits in memory formation. To test this hypothesis, we employed 18-month-old hAPP(J20) along with littermate controls. Animals of this age were utilized so as to ensure pronounced AD-like deficit at the start of the experiment.

Animals were evaluated for spatial learning and memory via a modified version the of Morris water maze. We employed a design similar to the design employed by Artinian and colleagues [39] where they performed familiarization of animals to the maze on day 1, acquisition training on day 2 followed by injection of a proteasome inhibitor and a probe trial on day 3. In this design they reported treatment with the proteasome inhibitor to produce spatial memory deficits [39]. In our design, animals were first acclimated to the maze using a raised platform at multiple entry locations on day 1. On day 2 animals received an IP injection of TAT-DEN then 3 training sessions at multiple entry sites with the platform submerged. On day 3 a probe trial was performed. We demonstrated that hAPP(J20) mice show deficits in memory formation showing reduced latency to find the submerged platform in the final training session, while hAPP(J20) mice treated with TAT-DEN showed improvement in this measure. TAT1-DEN produced no effect on memory acquisition in control mice (**Fig. 7F**). Furthermore, we show TAT1-DEN to rescue deficits in memory retention. hAPP(J20) mice showed deficits during the next day probe trial with deficits in latency to reach the platform zone, in comparison to control animals. In contrast hAPP(J20) mice treated with TAT-DEN showed similar latency to the platform zone as control animals (**Fig. 7F**).

Our results demonstrate Aβ impacts on learning memory and cell death to be in part dependent on its role as a proteasome inhibitor and that proteasome activation can rescue AD-like cognitive deficits both over chronic and acute treatment.

## DISCUSSION

Our study shows that oligomeric Aβ42 impairs proteasome function by binding to the lateral side of the 20S core, altering gate dynamics and thus the substrate access, and simultaneously lowering the affinity of the 20S core to the 19S regulatory particle. Prior work has shown that aggregated Aβ can inhibit proteasome activity [1, 5–8], with proposed mechanisms including allosteric gate stabilization in the 20S core [11]. We advance this work by demonstrating that oligomeric Aβ42 hampers allosteric gate opening, favouring transitions between closed and intermediate-gate conformers and stabilizing the latter. This stabilization is prominent even in the presence of a model substrate, with clearly affected allosteric mechanism of the substrate uptake. Importantly, impairing dynamics of the gate and the α face also compromises the 20S-19S interactions and shifts the equilibrium toward the free 20S particles rather than the 26S assembly, single or double-capped. These effects are specific to the oligomeric form of Aβ42 and are not observed with monomeric or fibrillar species. These findings extend prior studies by linking Aβ42 oligomer binding to the loss of structural and functional plasticity of the proteasome. Moreover, we propose the model of lateral rather than gate-area binding of the oligomers, with the functional defects resulting from the shifts in the conformational ensemble rather than direct blocking the entry of substrates. Such model is also consistent with the observed destabilization of the 26S proteasome assembly. Likely oligomers’ low binding affinity to 26S restricted our capability to detect decoration with Aβ with native electrophoresis. Together, our model offers a mechanistic framework for proteasome dysfunction in Alzheimer’s disease.

Pharmacologic activation of the proteasome with a small peptidomimetic TAT-DEN restores catalytic activity and conformational flexibility of the 20S proteasome in the presence of oligomeric Aβ42. We reported before that TAT-DEN and a homologous peptidomimetic TAT-TOD bind to the pocket between α1 and α2 subunits, activate the 20S and 26S proteasomes *in vitro* and *in vivo*, and rescue AD-like deficits in animal models [1, 27]. Now, we reinforce the notion of α1- α2 binding site for TAT-DEN ligand and suggest a specific “shaking off” the Aβ42 oligomers from the core, particularly when it assumes the intermediate – gate conformation. We propose that TAT-DEN binding to the α face allosterically lowers the affinity between the 20S core and the oligomers thus both protecting against and rescuing from the deleterious effects of Aβ42 *in vitro*, *in cellulo*, *and in vivo*.

Indeed, in cellular assays, proteasome treatment with 1μM TAT-DEN reduced Aβ-induced toxicity, while *in vivo* treatment protected against memory and survival deficits in *Drosophila* and improved cognitive performance in transgenic AD mice. Prior work demonstrated that proteasome activation reduces Aβ production by modulating precursor processing [1]; the current findings show that correcting proteasome dysfunction downstream of aggregation is sufficient to block Aβ-induced toxicity, even when initiated after pathological insult. This establishes a mechanistic link between proteasome restoration and functional rescue in models of Alzheimer’s disease.

Our observations stand in contrast to a prior study, which found Aβ inside the proteasome channel using cryoEM [41]. Instead, our findings show Aβ oligomers bound externally to the proteasome’s lateral surface, potentially reflecting differences in the species and structure of Aβ used. Our findings do not contradict the cryoEM data. Whereas the prior study used monomeric Aβ40, we used larger oligomeric Aβ42, expected to interact differently with the proteasome. It is worth noting that the proteasome-interacting oligomers that we detect are likely relatively small. Low molecular weight Aβ42 oligomers are the most toxic [42] and sufficiently structurally flexible [43] to coat the proteasome with relatively modest perturbations of morphometric parameters.

In our previous and current studies we demonstrated impairment of both 20S and 26S proteasomes function under AD, with the rescue provided by the employed proteasome agonists. It is presently unclear if AD – related defects are promoted by dysfunction of the 20S proteasome (ubiquitin-independent degradation), 26S proteasome (mostly ubiquitin-dependent degradation) or impact both. Considering the widespread involvement of both ubiquitin dependent and independent proteolysis in the neural proteostasis it is plausible that both proteasome species play distinct but critical roles. The prevailing understanding for how proteasome dysfunction moderates cognitive deficits is through programmed control of such pathways as the CREB transcription factors modulating long term potentiation [12–20], and modulation of the NMDA receptor impacting growth of dendritic spines [21–25]. These functions depend on turnover of polyubiquitinated proteins by the 26S proteasome, not the 20S proteasome. In contrast, emerging data suggests novel roles for 20S proteasome. Moreover, there is accumulating evidence of the presence of a novel plasma membrane proteasome in neurons. The authors report that approximately 40% of proteasome (specifically 20S proteasome) in neurons have a plasma membrane localization, which was not observed in non-neuronal cell cultures [44, 45]. Compromising these plasma membrane proteasomes impairs calcium signalling, thus affecting the key neuronal processes in neuronal function [45, 46]. An interesting notion open for further exploration is that the plasma membrane proteasomes may be the major species driving the vicious cycle of the formation of toxic Aβ oligomers, proteasome inhibition and the loss of proteostasis. Accordingly, the plasma membrane proteasomes may be the target of beneficial augmentation.

In our prior studies we suggested that the protective impacts from proteasome augmentation stem from increased turnover of amyloid precursor machinery thus slowing AD progression [1]. In this study we employed models which involved either the established deficits or treatment with preformed amyloid. This was undertaken intentionally to separate impacts of amyloid precursor synthesis and amyloid formation from amyloid toxicity. Indeed, we demonstrated that protective effects still exist both under acute treatment in animals with pre-existing deficits and in models treated with pre-formed Aβ, expected to inhibit critical proteasome functions [47]. We hypothesize that proteasome-dependent loss of proteostasis in neurons contributes to AD pathology that can be mitigated or even reversed under treatment with proteasome agonists.

Notably, this investigation was focused on Aβ, however tau is also highly impactful to AD and to proteasome function, with reports of significant inhibition of the proteasome by the hyperphosphorylated tau [9]. Further investigation is needed to uncover the impacts of proteasome modulation on tauopathy and tau toxicity, and potential cross-talk between toxicities induced by both proteins.

In conclusion, our data uncovers a distinct mechanism by which Aβ42 oligomers decorate lateral surface of the 20S core and severely restrict the allosteric movements of the proteasome gate, the ability of the core to accept substrates, and to form the 26S assembly. We demonstrate that a specific allosteric ligand of the α face is able to decrease the binding affinity between the 20S and oligomers and rescue the proteasome’s catalytic process. We further show that the proteasome impairment is a downstream driver of AD cognitive defects, whereas the proteasome activation is sufficient to prevent AD-like cognitive deficits.

## METHODS

### Key reagents

Purified 20S Proteasome (R&D Systems, Cat# E-360), purified 26S Proteasome (R&D Systems, Cat# E-365).

### Preparation of Aβ

Purified Aβ42 (Genscript, Cat# RP20527-1), was solubilized in 1% NH_3_ at 12.5mg/ml then diluted to 1mg/ml in PBS. For monomers, samples were sonicated for 60 minutes, vigorously vortexed for 1 minute then centrifuged at 16,000g for 10 minutes to remove aggregates. For oligomers, samples were sonicated for 60 minutes, incubated at 4°C for 24hr under gently agitation, vigorously vortexed for 1 minute then centrifuged at 14,000g for 10 minutes to remove aggregates. For fibrils, samples were sonicated for 60 minutes, vigorously vortexed for 1 minute then incubated at 37°C for 24hr. Consistent with prior studies [34]. The size distribution of the oligomers in the supernatant was examined by atomic force microscopy (AFM).

### Proteasome activity assay

The chymotrypsin-like activity was tested with 50 μM Suc-LLVY-AMC (succinyl-LeuLeuValTyr- 7-amino-4-methylcoumarin) model substrate, in a 96-well plate format, as previously described [1]. The reaction buffer consisted of 50 mM Tris-HCl (pH 7.8) + 100mM KCl. For testing the activity of the 26S holoenzyme, the 50 mM Tris-HCl buffer was supplemented with “MAD”: 1 mM MgCl_2_, 2 mM ATP, and 1 mM DTT (dithiothreitol).

### Atomic Force Microscopy (AFM) Imaging

Monomers, oligomers and fibrils of Aβ were imaged using the tapping (oscillating) mode in air with TESP probes and a scanner E of the Multimode Nanoscope IIIa (all from Bruker Inc., Santa Barbara, CA), as previously described for tau [48]. Live 20S or 26S proteasome particles were imaged with the tapping mode in liquid using our established procedure (31, 35), with SNL (Sharp Nitride Lever; Bruker) probes. Samples of proteasomes were mixed with Aβ oligomers (2 µM) and/or TAT activator (1 µM), or the respective vehicles (0.008% ammonia water for oligomers, 1% DMSO for TAT). The mixes were incubated for 1 hour at room temperature before (i) testing the chymotrypsin-like activity as above and (ii) diluting in “imaging buffer” (5 mM Tris-HCl, pH 7) and depositing on muscovite mica for electrostatic attachment. 26S proteasome preparations were incubated with the vehicle or oligomers for 1 hour as well, in the presence of “MAD”. Under these wet imaging conditions none of the Aβ species attached to the mica, leaving only proteasomes available for imaging. When indicated, 20S proteasomes were deposited on mica first, scanned, then treated on mica with Aβ oligomers (“Aβ pre-decorated”) and then after at least 30 min of incubation and scanning, treated with 1 µM TAT-DEN. Scanning and imaging was performed in height mode, with trace (left to right) and retrace (right to left) images collected, as described (31). The images of 1 µm^2^ fields (512 × 512 pixels) typically contained from 20 to 200 top-view (“standing”) 20S proteasome particles, with the top of α ring (“α face”) covered by a six-pixel scan line fragment and amenable for analysis. Only the standard 1^st^ order flattening (Nanoscope Analysis 1.70 software) or line-wise levelling (SPIP 6.013 software; Image Metrology, Denmark) was applied before analysis so the images should be considered “raw”. The raw numerical values of pixels of scans across α faces collected with a practical vertical resolution reaching 1 Å were used to discern between the three conformational forms: closed, intermediate and open, depending on the presence of a “dip” (local minimum; open form), a concave function (intermediate form) or a convex function (closed form) [49, 50]. Selected fields were scanned multiple times to follow conformational changes in the same set of particles. For the morphometric analysis of 20S or 26S particles the Particle Analysis function in SPIP software was used [51]. Frequency analysis, peak fitting and statistical analysis was performed with the OriginPro 2019 (OriginLabs, Northampton, MA).

### In silico docking of TAT-DEN

To identify molecular interactions of TAT-DEN with 20S proteasome, we performed molecular dynamics simulations of the ligand molecule followed by its docking studies. To address conformational flexibility of TAT-DEN, we applied temperature replica exchange molecular dynamics (T-REMD) [52]. The identified centroid conformations [53] were docked individually to control human 20S proteasome using molecular docking software Rhodium™ (Southwest Research Institute, San Antonio, TX). The scoring metrics were built from cavity occupancy (CAVOC, ligand selectivity for the binding pocket) and pose population scores (POP, binding pocket location). Based on the docking results, we identified TAT-DEN poses that bind in the α1-α2 grove with the high POP score and the highest CAVOC score [54]. This binding location for TAT-TOD was also identified previously by biochemical and *in silico* experiments [27].

### Patient samples

Flash-frozen hippocampal tissue was provided by the NIH NeuroBioank. Tissues were provided by multiple repositories; thus, some sites provided brain slices, while others provided powderised tissues. Samples were requested to have a post-mortem interval of less than 24 hours, from patients between 50 and 80 years old. Samples were powderised if needed and the materials weighed. For proteasome activity assays, samples were resuspended and homogenized in proteasome activity buffer. Data shown represents reanalyzed data from our prior publication [1].

### Olfactory aversion training

Experiments were performed as described in [55]. Animals were exposed (via an air pump) in alternation to two neutral odors (3-octanol and 4-methylcyclohexanol, prepared as a 1:10 dilution in mineral oil) for 5 min under low red light, and a 100-V 60-Hz shock was applied during exposure to one of the two odors. The odor associated with the electric shock was alternated between vials. After three training rounds per odor, animals were given 1 hour to recover then placed in a T maze (CelExplorer Labs) with opposing odors from either side. Flies were allowed 2 min to explore the maze, after which the maze sections were sealed, and the number of flies in each chamber was scored.

### Cell viability

Cells were maintained in clear 96-well plates. On the day of assay, 10 μl of WST-1 reagent (11644807001, Sigma-Aldrich) was added to each well, and cells were incubated in a 37°C, 5% CO2 incubator for 2 to 4 hours. Absorbance was measured at 450 nm using a Gemini series spectrophotometer.

### Osmotic pump implantation

Adapted from [56], animals were anesthetized with vaporized halothane, and a micro-osmotic pump (Alzet #2004) was stereotaxically implanted into the right lateral cerebral ventricle (at coordinates−1.0 mm mediolateral and −0.5 mm anterioposterior from Bregma;−1.5 mm dorsal-ventral from skull). Pumps contained either Aβ1-42, Aβ1-42 + TAT-TOD, Aβ1-42 + TAT-DEN or vehicle alone. Pumps were placed under the skin and wounds sutured closed. The pump has an expected release rate of 0.25 µl/hr for 4 weeks. Pumps were loaded with 55µg Aβ42 for an expected release of 68ng/h and 9.9mg/100ul of TAT-TOD/DEN to produce an anticipated tissue concentration of 100nM.

### Morris water maze

This test provides measures of hippocampal-dependent spatial learning and memory. A 121-cm water maze was used. Animals were given a series of three trials, ∼30 min apart, per day to find a submerged platform (∼1 cm below water level) in a large tank filled with water made opaque through the addition of white tempera-based nontoxic paint at 23.0° ± 1.0°C, in a room separated from the operator by a curtain. The pool was surrounded by large panels with geometric black and white designs that serve as distal cues. Maximum trial time was limited to 60 s, whereupon mice were guided to the platform. Mice were allowed to remain on the platform for 5 s and then were gently towel-dried and moved to their home cage under a heating lamp until dry. At the end of training, a probe trial was conducted where the platform was unavailable to measure the retention of the former platform location. The time each animal spent in the quadrant formerly containing the platform and the number of passes over that location provided a measure of memory. Data were collected using TopScan (CleverSys) by operators blinded to genotype.

### Y maze

Working memory is assessed by placing animals in a Y-shaped maze made of white Plexiglas with three arms, with equal angles between all arms. Each animal is placed in an arm of the maze and allowed to move freely around the apparatus, while the sequence and number of arm entries for each animal during a 5-min period are recorded manually by an experimenter blinded to the genotype. The number of spontaneous alternations, which occur when a mouse enters a different arm of the maze in each of three consecutive arm entries (e.g., ABC, CAB, or BCA but not BAB), was counted, and percent alternation was calculated as: (# of spontaneous alternations)/(total arm entries − 2) × 100.

### Study approval

All mouse studies performed were approved by the Institutional Animal Care and Use Committee at the University of Alabama at Birmingham (protocol 22179; Pickering, PI).

## ACKNOWLEDGMENTS

This work was supported by the National Institute of Aging R01 AG065301 (to A.M.P.), National Institute of General Medical Science R01 GM069819 (to M.G.), William and Ella Owens Medical Research Foundation (to M.G. and P.A.O.), National Institute of Health S10 Shared Instrumentation Grant OD034419-01 (to P.A.O.) and Fulbright Scholarship program (MB).

---

**Supplemental Fig. 1.**
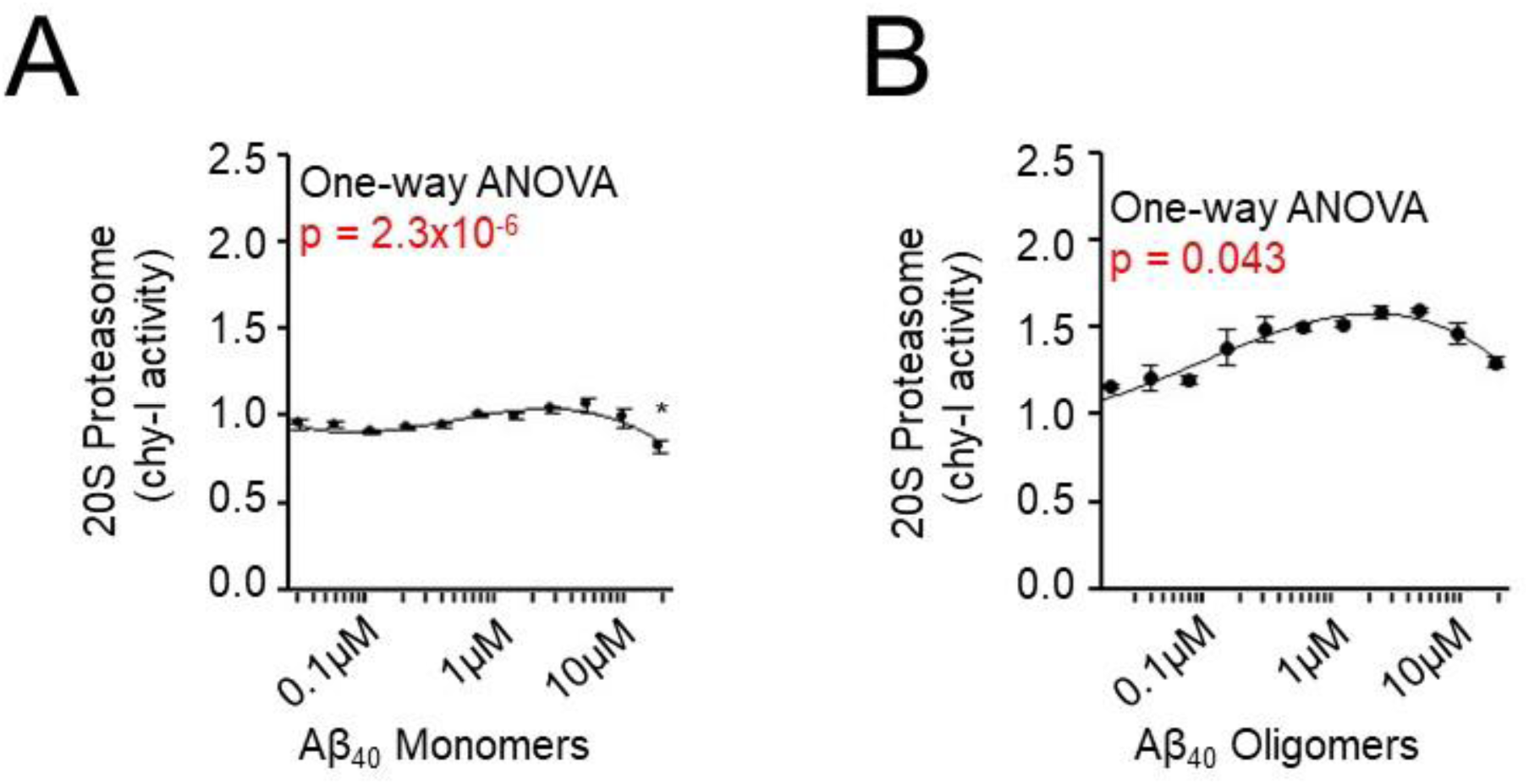
Aβ40 (unlike Aβ42) has minimal impacts on 20S proteasome activity. Chymotrypsin-like activity of purified human 20S proteasome after incubation with **(A)** monomeric Aβ40 or **(B)** oligomeric Aβ40. N=4 per group. Statistical comparison for performed by one-way ANOVA; asterisks indicate significance by Tukey’s post hoc test (p<0.05) with adjustment for multiple comparisons.

**Supplemental Fig. 2.**
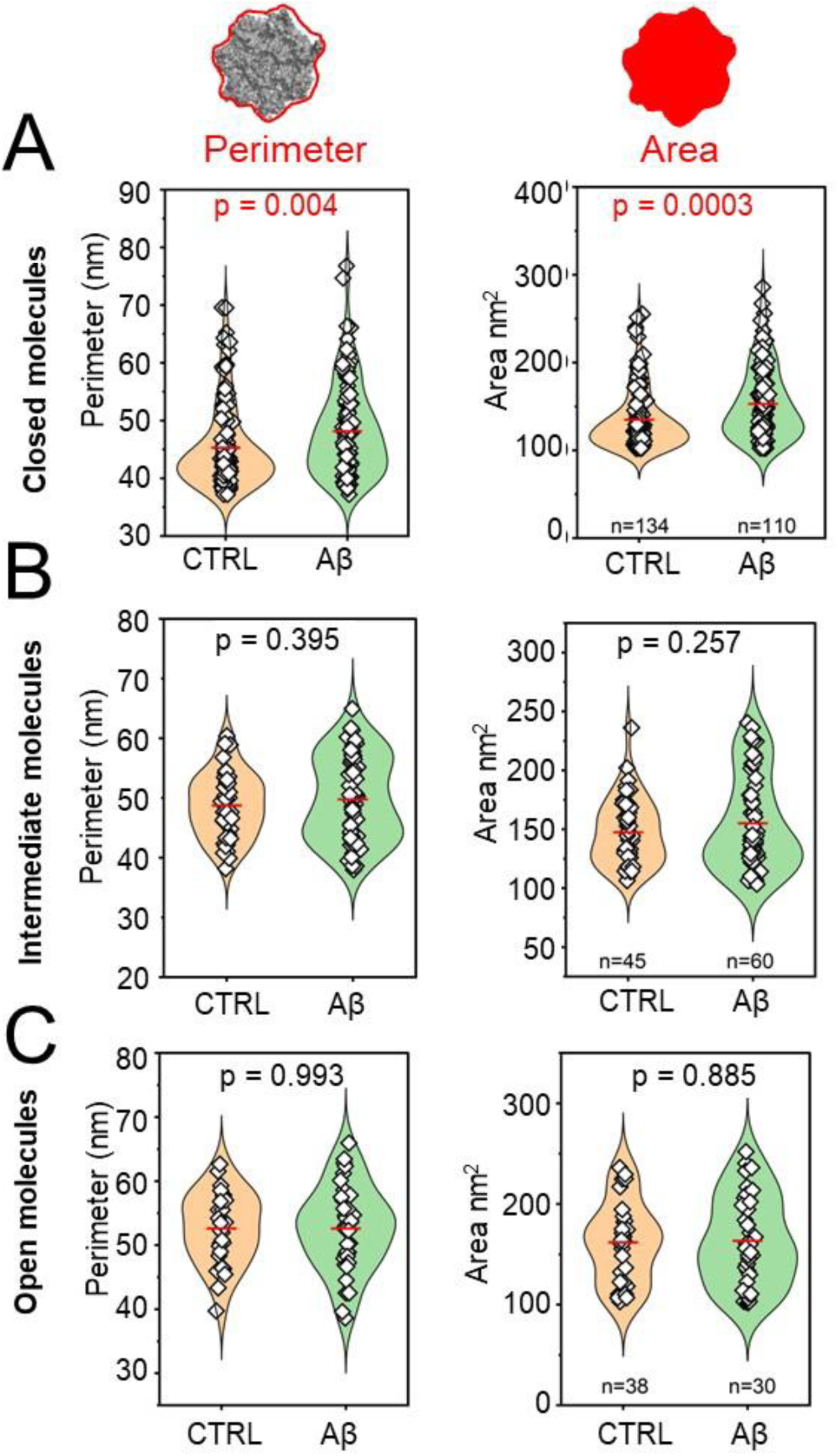
Treatment with oligomeric Aβ42 induces changes in shape of selected 20S proteasome conformers as shown with additional morphometric parameters. Morphometric analysis of top-view 20S proteasome particles imaged by AFM in the absence or presence of oligomeric Aβ42 and classified into three conformational states as in Fig. 2 was performed for two additional parameters: Perimeter (*Left*)– the total length of the projected contour, and Area (*Right*) – the projected footprint of a particle. Control group included 134 closed, 45 intermediate, and 30 open gate particles; Aβ42-treated group included 110 closed, 60 intermediate, and 38 open gate particles. Statistical analysis was conducted via pairwise comparison; p < 0.05 was considered significant.

**Supplemental Fig. 3.**
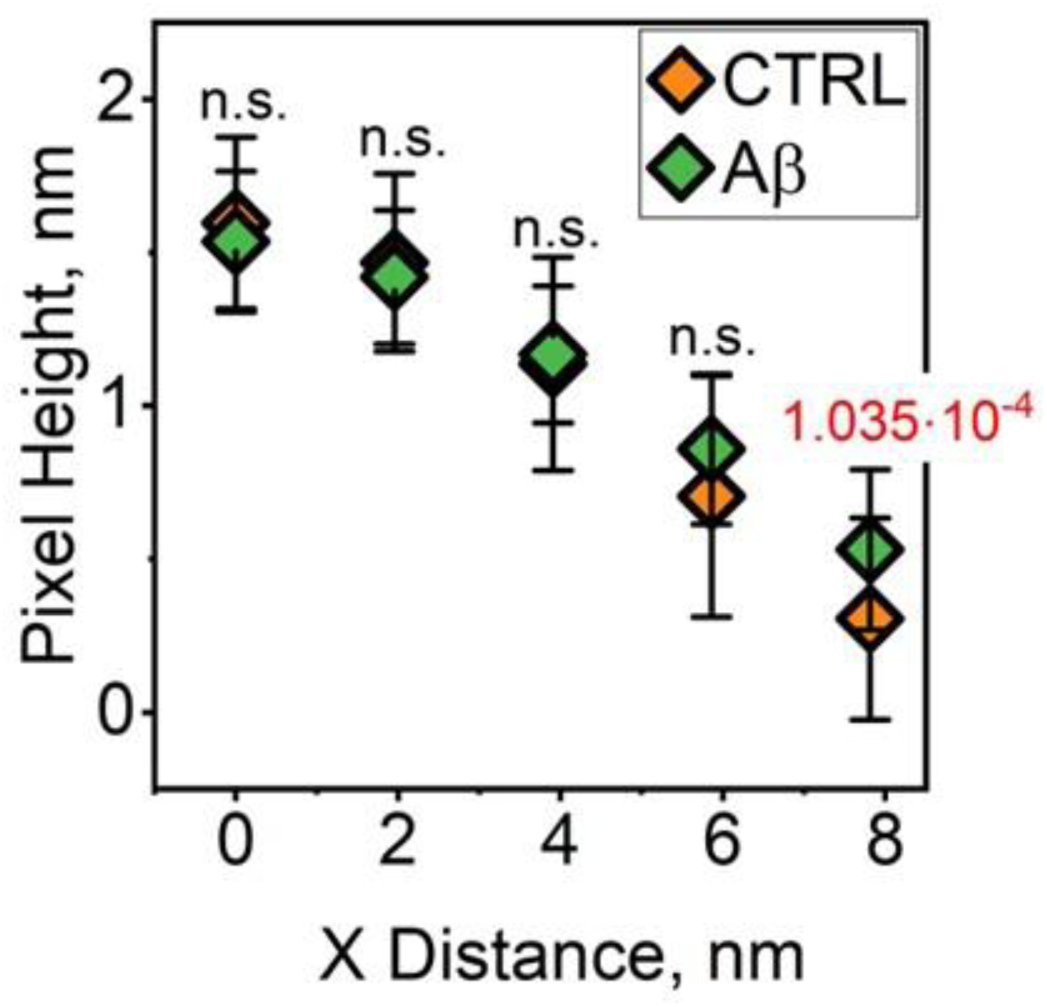
Morphometric properties of the top surface of the closed-gate proteasome α ring (α face) support a hypothesis of lateral rather than top-binding of the Aβ42 oligomers. We compared the raw Z values (height) of the pixels that form the center line of top view 20S proteasomes in the closed conformers of randomly selected 51 control and 70 oligomers-treated particles. Distance = 0 nm represents the center (gate area) of the particle. The α face radius extends to ∼ 5.5 nm (the determined diameter of α face in crystal structure or CryoEM models is ∼11 nm; for example: [3]. The height of pixels measured along the center line did not differ significantly between control and oligomers-treated particles, with no indication of an extra mass on a top of the treated particles. Instead, the significantly higher Z value was found only for a pixel beyond the expected diameter of the α face in oligomers-treated particles. The location of a taller pixel at the rim of the α ring is consistent with the lateral binding of oligomers, and the “broadening” of X extend (real time diameter) of Aβ42 oligomers- treated proteasomes.

**Supplemental Fig. 4.**
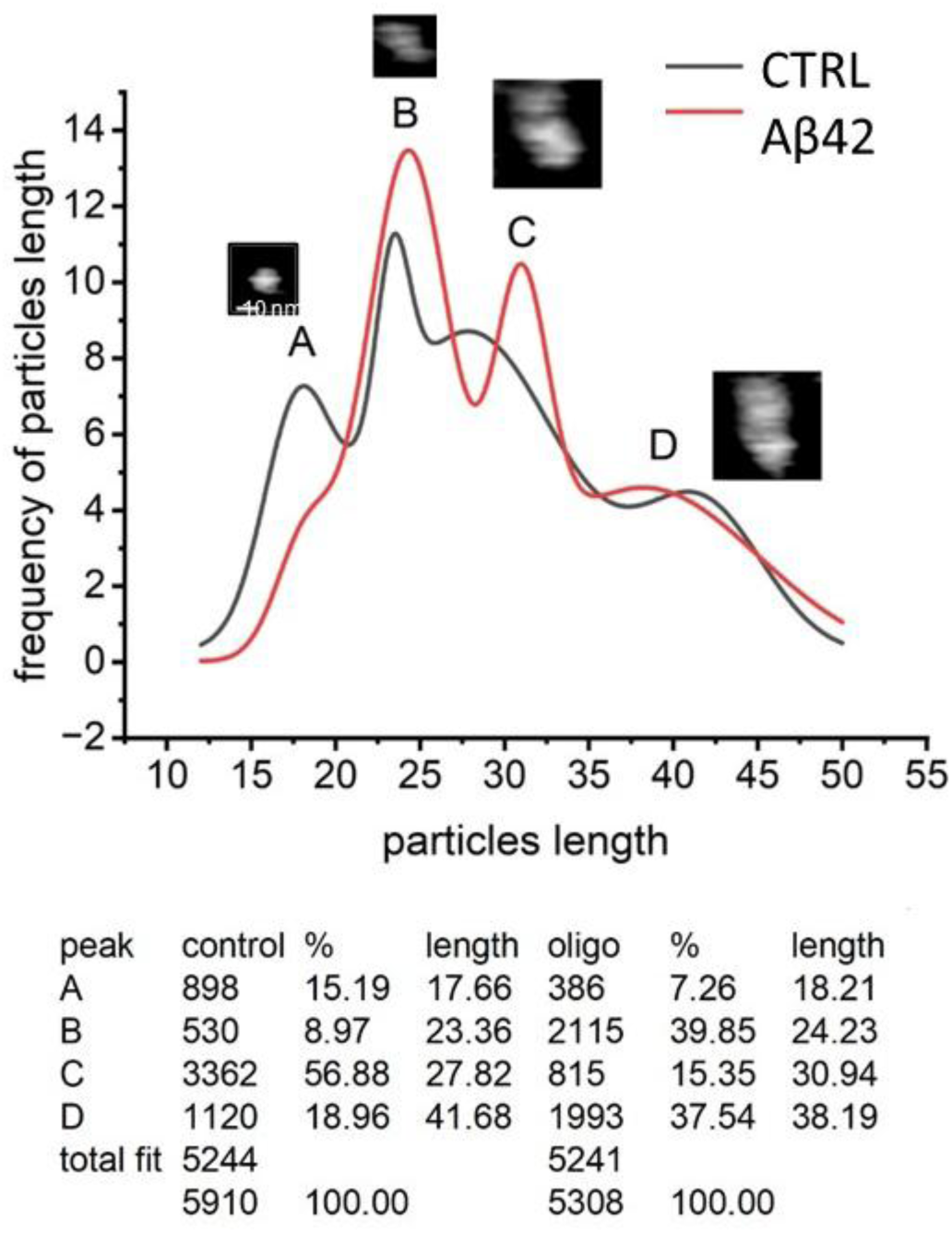
Frequency histogram of lengths of AFM-imaged 26S particles. The Length parameter (length of the particle’s oriented bounding rectangle parallel to the longest axis; a.k.a. maximal length of a particle) of 536 single control particles and 382 Aβ42 oligo-treated particles was analyzed. The 26S particles were imaged side-view and represent the longest objects: peaks C and D correspond roughly to single and double-capped proteasomes. Peaks A and B, clearly discerned in control samples, correspond to free top-view core particles and free 19S caps. Importantly, peak B was overrepresented in oligo-treated preparations, whereas peaks C and D (capped cores) were underrepresented. The low content of particles in peak A (free top view cores) in the treated preparations can be explained by the additional contribution of enlarged, oligomers-coated cores to peak B. The columns in Fig. 4F show combined partitions of particles in peaks A+B (free cores and caps) and C+D (capped cores, capable of processing polyubiquitinated substrates).

**Supplemental Fig. 5.**
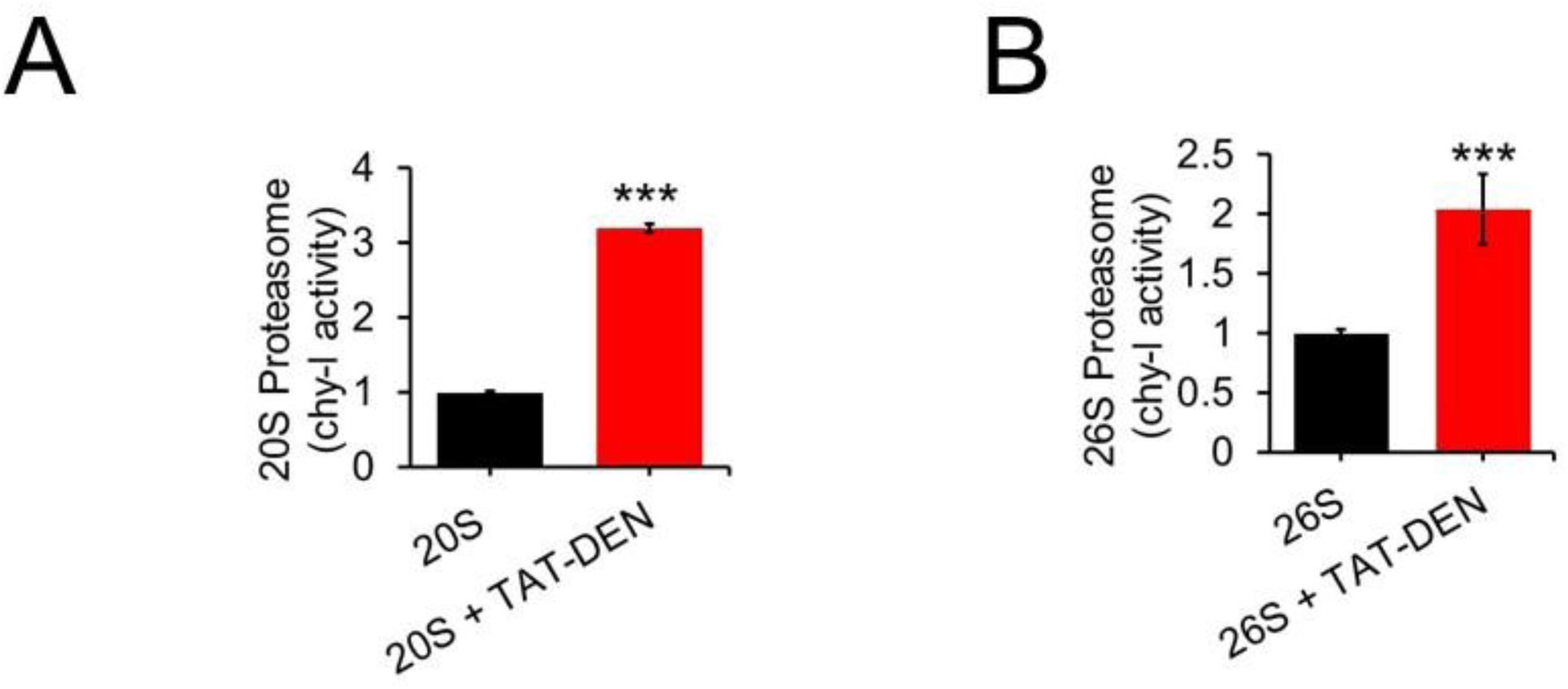
Peptidomimetic TAT-DEN activates the chymotrypsin-like activity of human proteasome. **(A)** purified human housekeeping 20S proteasome and **(B)** purified human 26S proteasome were incubated with 50 nM TAT-DEN. Model fluorogenic peptide Suc-LLVY-AMC was used as the substrate. N=4. Statistical analysis based on Student t-test ***P<0.001.

**Supplemental Fig. 6.**
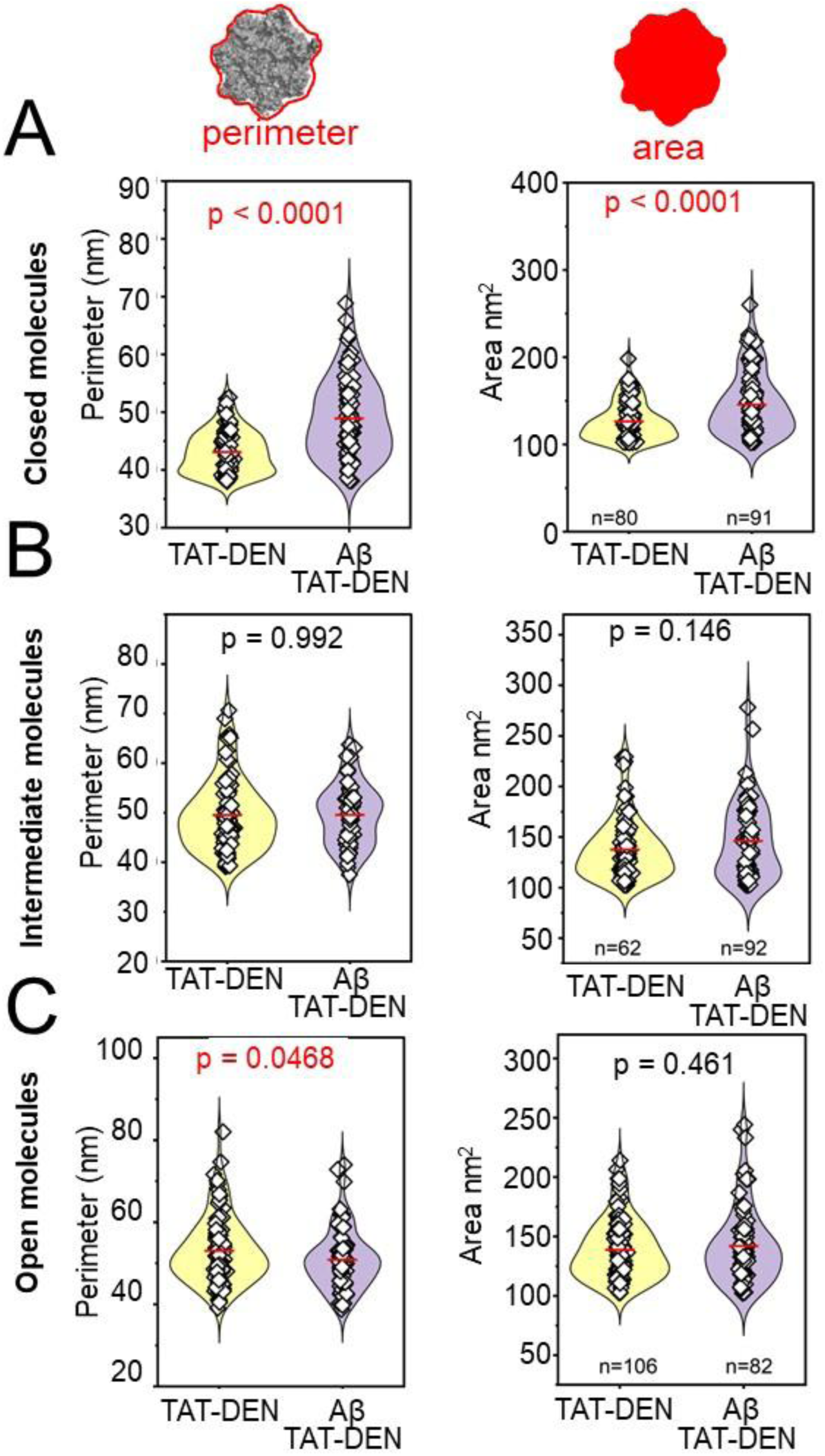
Morphometric shifts induced by TAT-DEN in 20S proteasome treated with oligomeric Aβ42 are limited to closed-gate conformers – analysis of additional morphometric parameters. Morphometric analysis of top-mounted 20S proteasome particles with AFM (tapping mode in liquid). Particles were classified into closed **(A)**, intermediate **(B)**, and open gate **(C)** conformers in the presence or absence of 2 μM oligomeric Aβ42 assessing: Perimeter (***Left***)– the total length of the projected contour, Area (***Middle***) – the projected footprint, Y-extent (***Right***) – the maximal length of particle along the Y axis. **(D)** In the proposed model closed conformers are laterally coated with Aβ oligomers, whereas intermediate conformers exhibit limited Aβ coating. Open conformers do not display coating by Aβ42 oligomers. Each point represents a single proteasome particle. TAT-DEN group included 80 closed, 62 intermediate, and 106 open gate particles; Aβ42 TAT-DEN - treated group included 91 closed, 92 intermediate, and 82 open gate particles. Statistical analysis was conducted via pairwise comparison; p < 0.05 was considered significant.

**Supplemental Fig. 7.**
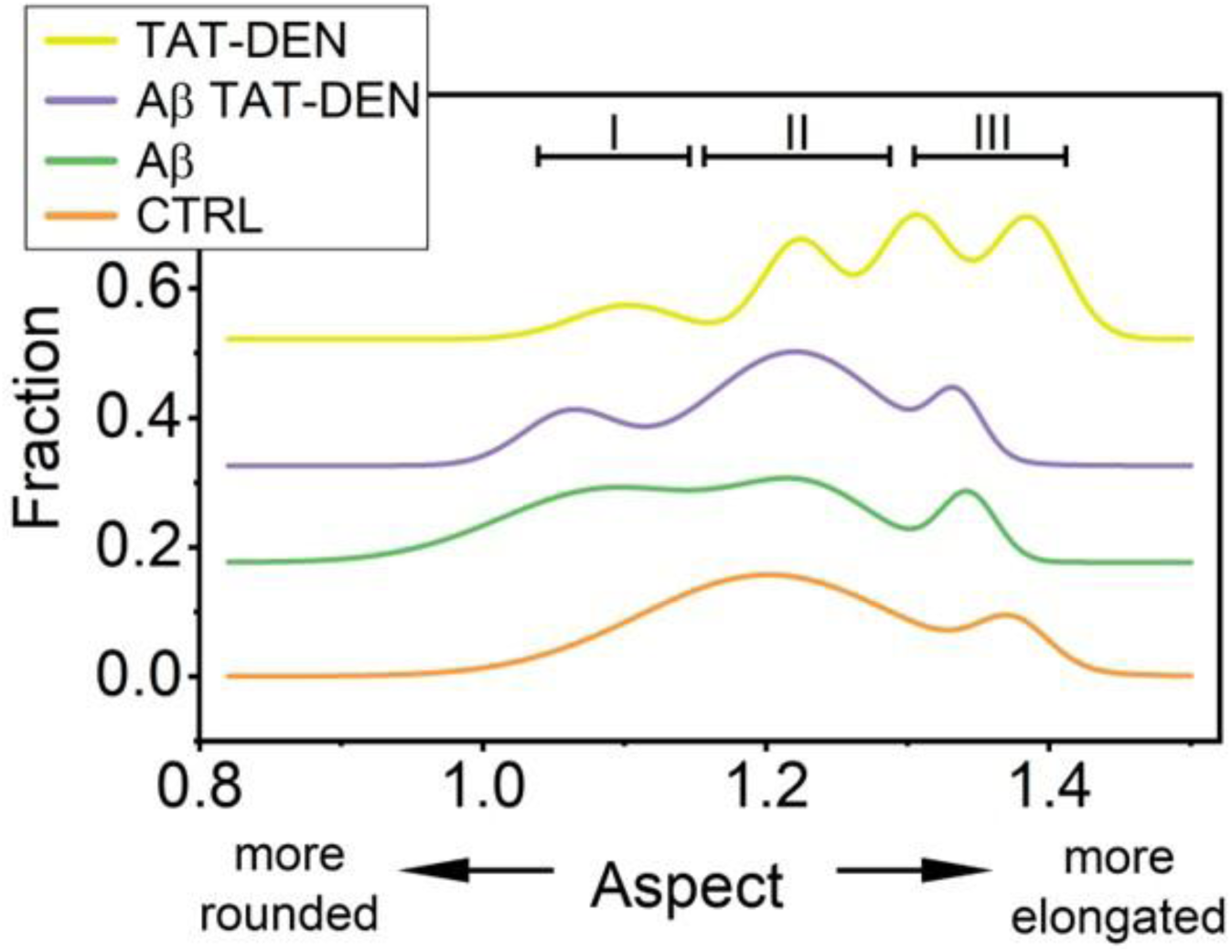
Aspect distribution in closed 20S particles depends on the bound ligand. Aspect was implemented as a measure of particle roundness (the ratio between maximal and minimal diameters measured from the particle’s center of gravity), It was significantly lowered in particles with putative lateral binding of oligomers (see Fig. 2**, 5**). Aspect was collected for 134 control, 110 Aβ42-treated, 80 TAT-DEN, and 91 Aβ42 TAT-DEN particles. We hypothesize that the roughly discerned Peak I contained mostly oligomers-coated particles, Peak II contained stable, oligomers-free particles and Peak III represented dynamic particles in transition between allosteric forms. Of note is putative unique dynamics of TAT-DEN treated proteasomes.

**Supplemental Fig. 8.**
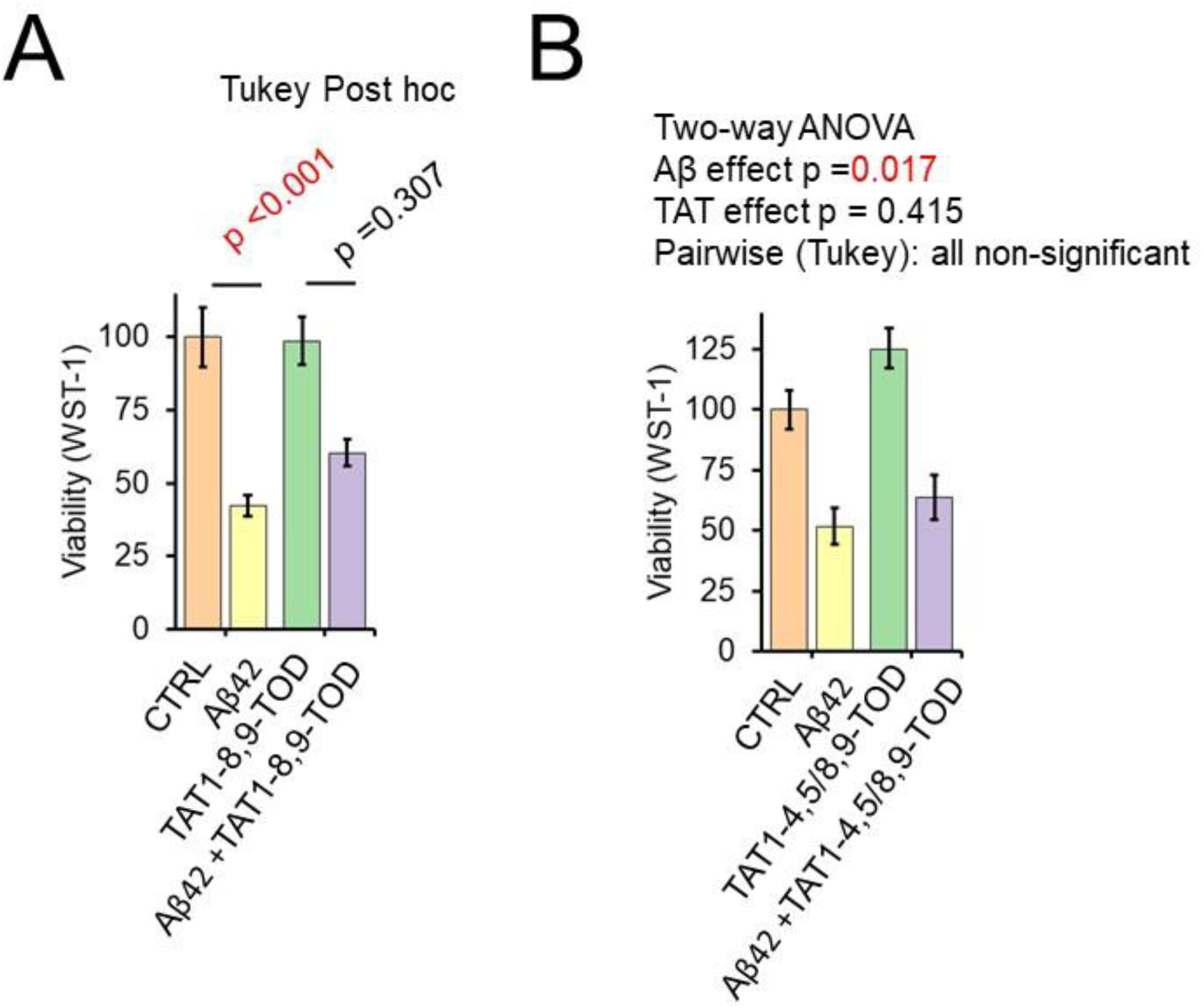
Aβ42 impairs cell viability. TAT-8,9TOD protects from Aβ42 induced cell death. A structurally similar TAT-4,5/8,9TOD, is a poor proteasome activator and does not protect from Aβ42. WST1, cell viability assays in SK-N-SH cells seeded at 100,000 cells per ml after 4 days treatment with 5 μM oligomeric Aβ42. Cell death was modulated under co-treatment with 1μM TAT-8,9TOD (A) or 1μM TAT4,5/8,9TOD (B). N = 15 wells/condition. Significance is based on two-way ANOVA with Tukey post hoc.

**Supplemental Fig. 9.**
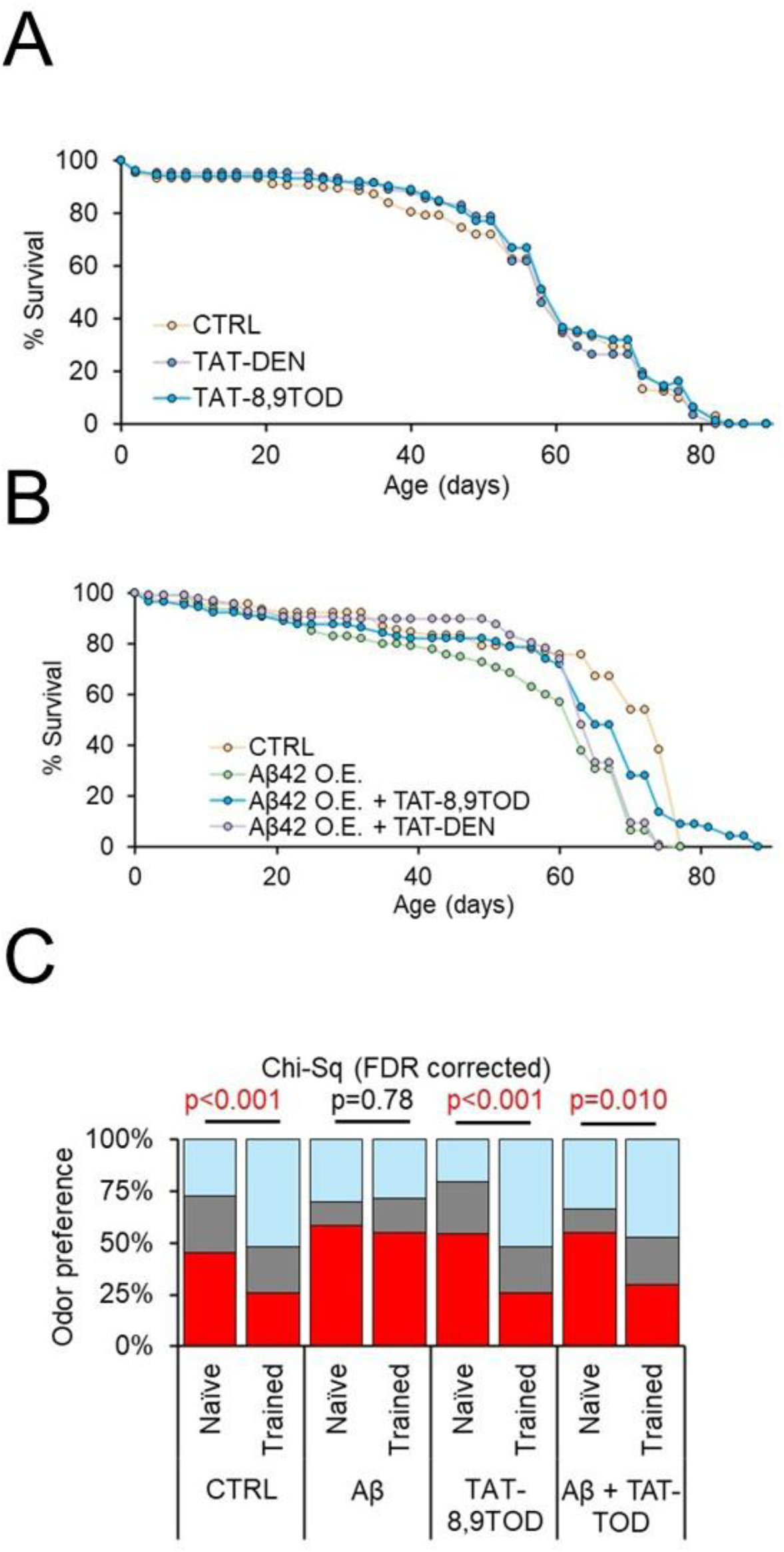
Proteasome augmentation rescues survival and cognitive deficits in a Drosophila Aβ42 overexpression model. **(A, B)** Survival assay in mated female Oregon-R flies **(A)** or Elav-GS-GAL4>UAS-Aβ42 flies **(B)** in the presence of 1μM TAT1-8,9TOD / TAT1-DEN or vehicle mixed directly into food. N = 89-150 animals/condition. **(C)** Olfaction aversion training assay in day 30 comparing Elav-GS-GAL4>UAS-Aβ42 ± 200 RU486 (CNT/Aβ42 O.E.) in the absence and presence of 1μM TAT1-8,9TOD or vehicle mixed directly into food. Significance of differences based on FDR corrected Chi-square analysis.

**Supplemental Fig. 10.**
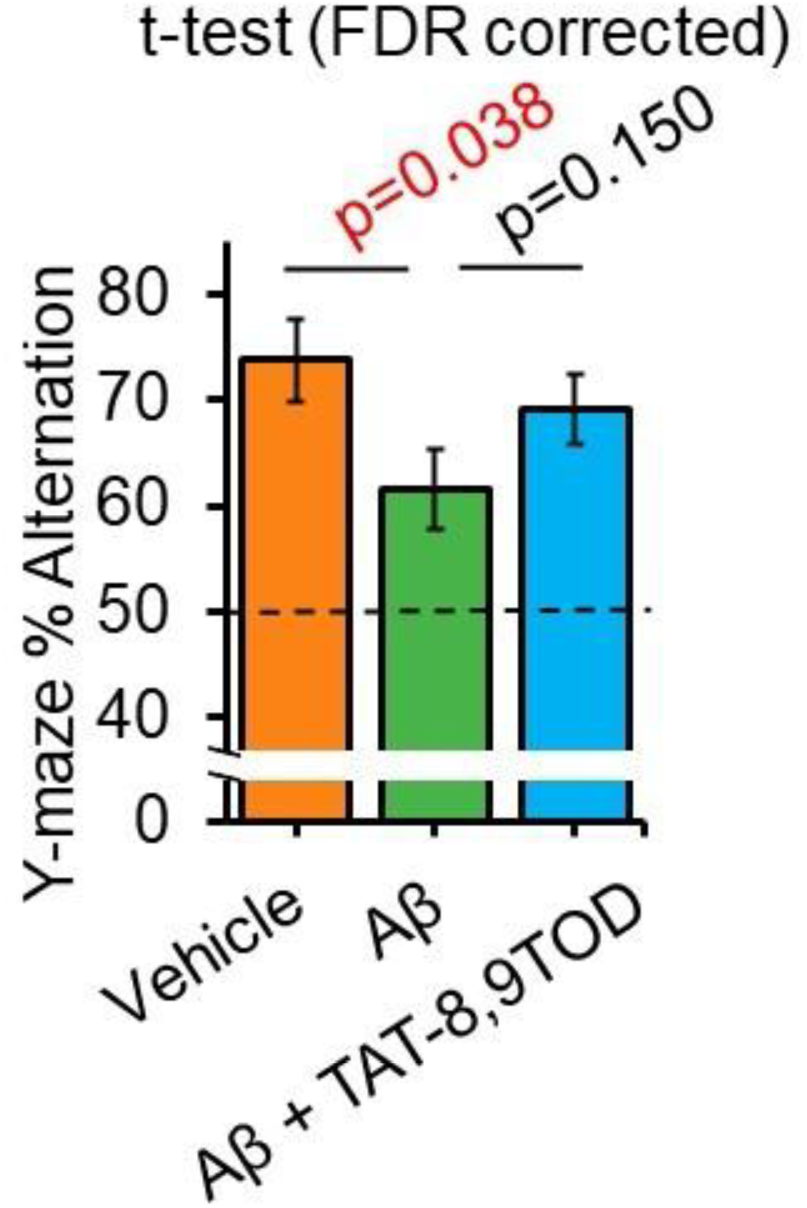
Aβ42 produces deficits in working memory Y-Maze. TAT-TOD produces a trend for improved performance. Osmotic pumps surgically implanted in 3 months old C57BL6J mice to deliver to the hippocampus +/- 1.25μmol of Aβ42 +/-100pmol of TAT-TOD per hr for 21 days. Spontaneous alternation in Y-Maze assessed. N=11 animals/group. Significance of differences based on FDR corrected T-test analysis.

**Supplemental Table 1.** Values of morphometric parameters and statistics for proteasome control particles and treated with Aβ42 oligomers. The data was used to generate violin plots in Fig. 2 and **Supplemental** Fig. 2.

|  | Area nm<sup>2</sup> |  | Z Max nm |  | Peri-meter nm |  | Aspect Ratio |  | X Extent nm |  |
| --- | --- | --- | --- | --- | --- | --- | --- | --- | --- | --- |
| **closed** | CTRL | Aβ | CTRL | Aβ | CTRL | Aβ | CTRL | Aβ | CTRL | Aβ |
| N total | 134 | 110 | 134 | 110 | 134 | 110 | 134 | 110 | 134 | 110 |
| Mean | 135.0 | 152.8 | 1.58 | 1.52 | 45.3 | 48.2 | 1.21 | 1.17 | 13.47 | 14.93 |
| Standard Deviation | 34.0 | 40.8 | 0.23 | 0.23 | 7.1 | 8.0 | 0.10 | 0.11 | 2.21 | 2.69 |
| Variance | 1157.7 | 1664.1 | 0.05 | 0.05 | 51.1 | 63.3 | 0.01 | 0.01 | 4.86 | 7.24 |
| Minimum | 102.1 | 102.5 | 1.10 | 1.06 | 37.2 | 37.2 | 0.90 | 0.86 | 9.78 | 11.09 |
| Median | 125.5 | 142.3 | 1.56 | 1.49 | 43.0 | 46.0 | 1.22 | 1.17 | 13.05 | 14.68 |
| Maximum | 255.3 | 285.9 | 2.60 | 2.22 | 69.6 | 76.8 | 1.39 | 1.37 | 20.87 | 24.79 |
| **Intermediate** |  |  |  |  |  |  |  |  |  |  |
| N total | 45 | 60 | 45 | 60 | 45 | 60 | 45 | 60 | 45 | 60 |
| Mean | 147.4 | 155.1 | 1.55 | 1.47 | 48.8 | 49.8 | 1.23 | 1.19 | 14.26 | 15.38 |
| Standard Deviation | 28.1 | 40.9 | 0.23 | 0.19 | 5.4 | 7.1 | 0.12 | 0.13 | 1.83 | 2.36 |
| Variance | 791.3 | 1670.2 | 0.05 | 0.04 | 29.0 | 50.0 | 0.02 | 0.02 | 3.36 | 5.56 |
| Minimum | 106.4 | 103.4 | 1.08 | 1.12 | 38.2 | 38.0 | 0.90 | 0.94 | 11.09 | 11.09 |
| Median | 144.2 | 141.1 | 1.56 | 1.44 | 48.9 | 48.9 | 1.25 | 1.17 | 14.35 | 15.00 |
| Maximum | 236.2 | 240.4 | 2.06 | 1.95 | 60.4 | 64.9 | 1.40 | 1.40 | 19.57 | 20.87 |
| **open** |  |  |  |  |  |  |  |  |  |  |
| N total | 30 | 38 | 30 | 38 | 30 | 38 | 30 | 38 | 30 | 38 |
| Mean | 161.9 | 163.4 | 1.44 | 1.43 | 52.6 | 52.6 | 1.25 | 1.20 | 16.26 | 16.29 |
| Standard Deviation | 39.7 | 44.1 | 0.23 | 0.22 | 5.5 | 6.6 | 0.09 | 0.13 | 2.03 | 2.14 |
| Variance | 1577.5 | 1940.5 | 0.05 | 0.05 | 30.8 | 43.0 | 0.01 | 0.02 | 4.11 | 4.57 |
| Minimum | 103.0 | 101.7 | 1.16 | 0.96 | 39.7 | 38.6 | 1.07 | 0.94 | 12.39 | 11.74 |
| Median | 163.8 | 158.1 | 1.38 | 1.41 | 52.9 | 52.9 | 1.25 | 1.20 | 16.63 | 16.31 |
| Maximum | 236.2 | 251.9 | 2.08 | 1.92 | 62.6 | 65.9 | 1.38 | 1.39 | 19.57 | 20.87 |

**Supplemental Table 2:**
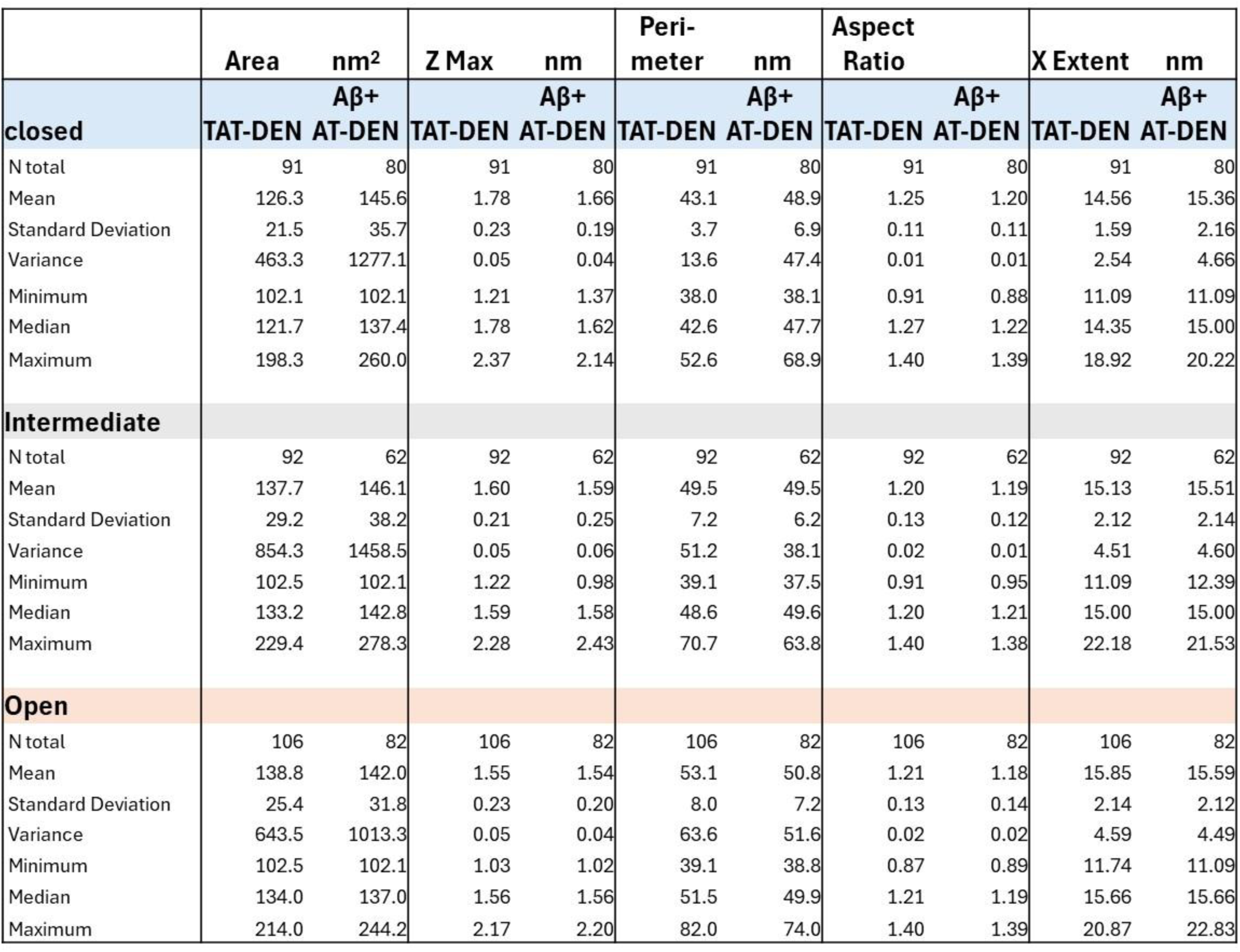
Values of morphometric parameters and statistics for proteasome particles treated with TAT-DEN or with Aβ42 oligomers and TAT-DEN. The data was used to generate violin plots in Fig. 5 and **Supplemental** Fig. 6.

## Bibliography

1. Chocron, E.S., et al., Genetic and pharmacologic proteasome augmentation ameliorates Alzheimer’s-like pathology in mouse and fly APP overexpression models. Sci Adv, 2022. 8(23): p. eabk2252.

2. Schrader, J., et al., The inhibition mechanism of human 20S proteasomes enables next-generation inhibitor design. Science, 2016. 353(6299): p. 594–8.

3. Dong, Y., et al., Cryo-EM structures and dynamics of substrate-engaged human 26S proteasome. Nature, 2019. 565(7737): p. 49–55.

4. Chen, S., et al., Structural basis for dynamic regulation of the human 26S proteasome. Proc Natl Acad Sci U S A, 2016. 113(46): p. 12991–12996.

5. Keller, J.N., K.B. Hanni, and W.R. Markesbery, Impaired proteasome function in Alzheimer’s disease. J Neurochem, 2000. 75(1): p. 436–9.

6. Rosen, K.M., et al., Parkin reverses intracellular beta-amyloid accumulation and its negative effects on proteasome function. J Neurosci Res, 2010. 88(1): p. 167–78.

7. Almeida, C.G., R.H. Takahashi, and G.K. Gouras, Beta-amyloid accumulation impairs multivesicular body sorting by inhibiting the ubiquitin-proteasome system. J Neurosci, 2006. 26(16): p. 4277–88.

8. Shringarpure, R., et al., 4-Hydroxynonenal-modified amyloid-beta peptide inhibits the proteasome: possible importance in Alzheimer’s disease. Cell Mol Life Sci, 2000. 57(12): p. 1802–9.

9. Keck, S., et al., Proteasome inhibition by paired helical filament-tau in brains of patients with Alzheimer’s disease. J Neurochem, 2003. 85(1): p. 115–22.

10. Myeku, N., et al., Tau-driven 26S proteasome impairment and cognitive dysfunction can be prevented early in disease by activating cAMP-PKA signaling. Nat Med, 2016. 22(1): p. 46–53.

11. Thibaudeau, T.A., R.T. Anderson, and D.M. Smith, A common mechanism of proteasome impairment by neurodegenerative disease-associated oligomers. Nat Commun, 2018. 9(1): p. 1097.

12. Soberg, K. and B.S. Skalhegg, The Molecular Basis for Specificity at the Level of the Protein Kinase a Catalytic Subunit. Front Endocrinol (Lausanne), 2018. 9: p. 538.

13. Greenberg, S.M., et al., A molecular mechanism for long-term sensitization in Aplysia. Nature, 1987. 329(6134): p. 62–5.

14. Hegde, A.N., A.L. Goldberg, and J.H. Schwartz, Regulatory subunits of cAMP-dependent protein kinases are degraded after conjugation to ubiquitin: a molecular mechanism underlying long-term synaptic plasticity. Proc Natl Acad Sci U S A, 1993. 90(16): p. 7436–40.

15. Chain, D.G., et al., Mechanisms for generating the autonomous cAMP-dependent protein kinase required for long-term facilitation in Aplysia. Neuron, 1999. 22(1): p. 147–56.

16. Hegde, A.N., et al., Ubiquitin C-terminal hydrolase is an immediate-early gene essential for long-term facilitation in Aplysia. Cell, 1997. 89(1): p. 115–26.

17. Upadhya, S.C., T.K. Smith, and A.N. Hegde, Ubiquitin-proteasome-mediated CREB repressor degradation during induction of long-term facilitation. J Neurochem, 2004. 91(1): p. 210–9.

18. Dong, C., et al., Proteasome inhibition enhances the induction and impairs the maintenance of late-phase long-term potentiation. Learn Mem, 2008. 15(5): p. 335–47.

19. Hegde, A.N., The ubiquitin-proteasome pathway and synaptic plasticity. Learn Mem, 2010. 17(7): p. 314–27.

20. Ferreira, J.S., et al., Interplay between NMDA receptor dynamics and the synaptic proteasome. Eur J Neurosci, 2021. 54(6): p. 6000–6011.

21. Bingol, B. and E.M. Schuman, Activity-dependent dynamics and sequestration of proteasomes in dendritic spines. Nature, 2006. 441(7097): p. 1144–8.

22. Bingol, B., et al., Autophosphorylated CaMKIIalpha acts as a scaffold to recruit proteasomes to dendritic spines. Cell, 2010. 140(4): p. 567–78.

23. Shen, H., et al., NAC1 regulates the recruitment of the proteasome complex into dendritic spines. J Neurosci, 2007. 27(33): p. 8903–13.

24. Hamilton, A.M., et al., Activity-dependent growth of new dendritic spines is regulated by the proteasome. Neuron, 2012. 74(6): p. 1023–30.

25. Hamilton, A.M., et al., A dual role for the RhoGEF Ephexin5 in regulation of dendritic spine outgrowth. Mol Cell Neurosci, 2017. 80: p. 66–74.

26. Ribeiro, F.C., et al., Synaptic proteasome is inhibited in Alzheimer’s disease models and associates with memory impairment in mice. Commun Biol, 2023. 6(1): p. 1127.

27. Osmulski, P.A., et al., New Peptide-Based Pharmacophore Activates 20S Proteasome. Molecules, 2020. 25(6).

28. Chondrogianni, N., et al., 20S proteasome activation promotes life span extension and resistance to proteotoxicity in Caenorhabditis elegans. FASEB J, 2015. 29(2): p. 611–22.

29. Mladenovic Djordjevic, A.N., et al., Pharmacological intervention in a transgenic mouse model improves Alzheimer’s-associated pathological phenotype: Involvement of proteasome activation. Free Radic Biol Med, 2021. 162: p. 88–103.

30. Vasilopoulou, M.A., et al., Healthspan improvement and anti-aggregation effects induced by a marine-derived structural proteasome activator. Redox Biol, 2022. 56: p. 102462.

31. VerPlank, J.J.S., et al., cGMP via PKG activates 26S proteasomes and enhances degradation of proteins, including ones that cause neurodegenerative diseases. Proc Natl Acad Sci U S A, 2020. 117(25): p. 14220–14230.

32. Davies, K.J., Degradation of oxidized proteins by the 20S proteasome. Biochimie, 2001. 83(3-4): p. 301–10.

33. Fu, L., et al., Comparison of neurotoxicity of different aggregated forms of Abeta40, Abeta42 and Abeta43 in cell cultures. J Pept Sci, 2017. 23(3): p. 245–251.

34. Stine, W.B., Jr., et al., In vitro characterization of conditions for amyloid-beta peptide oligomerization and fibrillogenesis. J Biol Chem, 2003. 278(13): p. 11612–22.

35. Osmulski, P.A., M. Hochstrasser, and M. Gaczynska, A tetrahedral transition state at the active sites of the 20S proteasome is coupled to opening of the alpha-ring channel. Structure, 2009. 17(8): p. 1137–47.

36. Bedford, L., et al., Assembly, structure, and function of the 26S proteasome. Trends Cell Biol, 2010. 20(7): p. 391–401.

37. Grune, T., et al., HSP70 mediates dissociation and reassociation of the 26S proteasome during adaptation to oxidative stress. Free Radic Biol Med, 2011. 51(7): p. 1355–64.

38. Lopez-Salon, M., et al., The ubiquitin-proteasome cascade is required for mammalian long-term memory formation. Eur J Neurosci, 2001. 14(11): p. 1820–6.

39. Artinian, J., et al., Protein degradation, as with protein synthesis, is required during not only long-term spatial memory consolidation but also reconsolidation. Eur J Neurosci, 2008. 27(11): p. 3009–19.

40. Lee, S.H., et al., Synaptic protein degradation underlies destabilization of retrieved fear memory. Science, 2008. 319(5867): p. 1253–6.

41. Gregori, L., et al., Binding of amyloid beta protein to the 20 S proteasome. J Biol Chem, 1997. 272(1): p. 58–62.

42. Yang, T., et al., Large Soluble Oligomers of Amyloid beta-Protein from Alzheimer Brain Are Far Less Neuroactive Than the Smaller Oligomers to Which They Dissociate. J Neurosci, 2017. 37(1): p. 152–163.

43. Liang, R., et al., Amyloid-beta oligomers, curvilinear and annular assemblies, imaged by cryo-ET, cryo-EM, and AFM. Sci Adv, 2025. 11(35): p. eadx9030.

44. Ramachandran, K.V. and S.S. Margolis, A mammalian nervous-system-specific plasma membrane proteasome complex that modulates neuronal function. Nat Struct Mol Biol, 2017. 24(4): p. 419–430.

45. Ramachandran, K.V., et al., Activity-Dependent Degradation of the Nascentome by the Neuronal Membrane Proteasome. Mol Cell, 2018. 71(1): p. 169–177 e6.

46. He, H.Y., et al., Neuronal membrane proteasomes regulate neuronal circuit activity in vivo and are required for learning-induced behavioral plasticity. Proc Natl Acad Sci U S A, 2023. 120(3): p. e2216537120.

47. Davidson, K. and A.M. Pickering, The proteasome: A key modulator of nervous system function, brain aging, and neurodegenerative disease. Front Cell Dev Biol, 2023. 11: p. 1124907.

48. Hussong, S.A., et al., Soluble pathogenic tau enters brain vascular endothelial cells and drives cellular senescence and brain microvascular dysfunction in a mouse model of tauopathy. Nat Commun, 2023. 14(1): p. 2367.

49. Gizynska, M., et al., Proline- and Arginine-Rich Peptides as Flexible Allosteric Modulators of Human Proteasome Activity. J Med Chem, 2019. 62(1): p. 359–370.

50. Giletto, M.B., et al., Pipecolic esters as minimized templates for proteasome inhibition. Org Biomol Chem, 2019. 17(10): p. 2734–2746.

51. Rodriguez, K.A., et al., A cytosolic protein factor from the naked mole-rat activates proteasomes of other species and protects these from inhibition. Biochim Biophys Acta, 2014. 1842(11): p. 2060–72.

52. Liu, P., et al., Replica exchange with solute tempering: a method for sampling biological systems in explicit water. Proc Natl Acad Sci U S A, 2005. 102(39): p. 13749–54.

53. Hess, B., et al., GROMACS 4: Algorithms for Highly Efficient, Load-Balanced, and Scalable Molecular Simulation. J Chem Theory Comput, 2008. 4(3): p. 435–47.

54. Stank, A., et al., Protein Binding Pocket Dynamics. Acc Chem Res, 2016. 49(5): p. 809–15.

55. Munkacsy, E., et al., Neuronal-specific proteasome augmentation via Prosbeta5 overexpression extends lifespan and reduces age-related cognitive decline. Aging Cell, 2019. 18(5): p. e13005.

56. Craft, J.M., et al., Aminopyridazines inhibit beta-amyloid-induced glial activation and neuronal damage in vivo. Neurobiol Aging, 2004. 25(10): p. 1283–92.

